# Contrasting genomic consequences of anthropogenic reintroduction and natural recolonisation in high-arctic wild reindeer

**DOI:** 10.1101/2022.11.25.517957

**Authors:** Hamish A. Burnett, Vanessa C. Bieker, Mathilde Le Moullec, Bart Peeters, Jørgen Rosvold, Åshild Ønvik Pedersen, Love Dalén, Leif Egil Loe, Henrik Jensen, Brage B. Hansen, Michael D. Martin

**Author notes:** corresponding authors*, **Corresponding author contact:** Hamish A. Burnett Michael D. Martin. contributed equally (Brage B. Hansen, Michael D. Martin).

## Abstract

Anthropogenic reintroduction can supplement natural recolonisation in reestablishing a species’ distribution and abundance. However, both reintroductions and recolonisations can give rise to population bottlenecks that reduce genetic diversity and increase inbreeding, potentially causing accumulation of genetic load and reduced fitness. Most current populations of the endemic high-arctic Svalbard reindeer (*Rangifer tarandus platyrhynchus*) originate from recent reintroductions or recolonisations following regional extirpations due to past overharvesting. We investigated and compared the genomic consequences of these two paths to reestablishment using whole-genome shotgun sequencing of 100 Svalbard reindeer across their range. We found little admixture between reintroduced and natural populations. Two reintroduced populations, each founded by 12 individuals around four decades (i.e. 8 reindeer generations) ago, formed two distinct genetic clusters. Compared to the source population, these populations showed only small decreases in genome-wide heterozygosity and increases in inbreeding and lengths of runs of homozygosity. In contrast, the two naturally recolonised populations without admixture possessed much lower heterozygosity, higher inbreeding, and longer runs of homozygosity, possibly caused by serial population bottlenecks and/or fewer or more genetically related founders than in the reintroduction events. Naturally recolonised populations can thus be more vulnerable to the accumulation of genetic load than reintroduced populations. This suggests that in some organisms even small-scale reintroduction programs based on genetically diverse source populations can be more effective than natural recolonisation in establishing genetically diverse populations. These findings warrant particular attention in the conservation and management of populations and species threatened by habitat fragmentation and loss.

## Introduction

Species reintroductions are increasingly being used in ecological restoration and biodiversity conservation programmes (Armstrong & Seddon 2008; Weeks *et al*. 2011; Seddon *et al*. 2014). Most reintroductions involve translocation of a small number of individuals to establish new populations that may be geographically isolated from the species’ current range (Frankham 2010). Founding populations are often characterised by having a small effective population size, only a subset of the genetic variation that exists in their source populations, and limited or no gene-flow with other populations (Lynch & Gabriel 1990; Frankham 2010). An alternative to reintroduction by translocations are more passive measures that facilitate natural dispersal and recolonisation of species’ ranges (Scott *et al*. 2001). Natural recolonisation processes may differ from reintroductions in that they require some connectivity with other populations. They can therefore be very slow, even in highly mobile species (Larter *et al*. 2000; Hurford *et al*. 2006). Especially in fragmented habitats and in species with low dispersal rates, natural recolonisation (including recolonisation from reintroduced populations) may also involve one or multiple sequential population bottlenecks and relative isolation of recolonised populations (Clegg *et al*. 2002; Pruett & Winker 2005).

Due to the often small founder population size, both reintroduced and naturally recolonised populations may initially experience strong genetic drift and accumulate inbreeding because individuals are more likely to share common ancestors, which increases homozygosity (Nei *et al*. 1975; Allendorf 1986). The levels of genetic diversity in reintroduced and recolonised populations are however also affected by population growth rate, genetic structure, and immigration (Latch & Rhodes 2005; Biebach & Keller 2010, 2012). For example, inbreeding and genetic drift accumulate over generations at a rate that depends on the population size and the rate of immigration (Whitlock *et al*. 2000; Willi *et al*. 2013). Rapid population growth reduces the duration of a population bottleneck and the degree of genetic drift (Nei *et al*. 1975; Allendorf 1986). Immigration counteracts the loss of diversity due to drift by introducing unrelated individuals and novel genetic material that replenishes genetic variation and reduces inbreeding rates (Vucetich & Waite 2000; Latch & Rhodes 2005; Frankham *et al*. 2017). The accumulation of inbreeding and genetic drift can allow low frequency (partially) recessive deleterious alleles that are rarely homozygous (i.e. masked genetic load (Bertorelle *et al*. 2022)) to increase in frequency and even become fixed. Consequently, masked genetic load may be converted to realised genetic load (Wang *et al*. 1999; Bertorelle *et al*. 2022), which is expected to reduce fitness (i.e. inbreeding depression (Charlesworth & Willis 2009)). However, this process exposes deleterious variation to selection, potentially purging strongly deleterious recessive alleles and reducing the fitness consequences of future inbreeding (Hedrick & Garcia-Dorado 2016; Robinson *et al*. 2018). Genetic drift also reduces genetic diversity, including potentially adaptive genetic variation that may be important for evolutionary responses necessary to maintain fitness in changing environments (Frankham 2005; Kardos *et al*. 2021). Together, these genetic consequences can impact both the short- and long-term viability of populations (Frankham 2005; Weeks *et al*. 2011).

Consequently, a key goal in the management of reestablishing or fragmented populations is to maximise the genetic diversity and minimise drift and inbreeding (Frankham *et al*. 2017). Several studies have shown that genetic diversity in reintroduced and naturally recolonised populations is often higher in those that receive gene flow from other populations (Latch and Rhodes 2005; Biebach and Keller 2012; Malaney et al. 2018), originate from multiple source populations (Williams et al. 2000; Huff et al. 2010; Williams and Scribner 2010; Sasmal et al. 2013; Vasiljevic et al. 2022), and in reintroductions that use multiple translocations (Drauch & Rhodes 2007; Cullingham & Moehrenschlager 2013). Without such mitigating factors, erosion of genetic diversity and accumulation of inbreeding can occur due to isolation and/or slow population growth (Williams *et al*. 2002; Hundertmark & van Daele 2010), and may have detrimental population-level consequences for fitness-related traits (Wisely *et al*. 2008) and population growth rates (Bozzuto *et al*. 2019). Differences in population connectivity and demography between reintroductions and natural recolonisations could therefore result in differing genetic consequences for these two paths to population reestablishment. Reintroduced populations may be more isolated from their source than naturally recolonised populations. However, the sequential reestablishment of habitat that often characterises natural recolonisation can result in cumulative founder effects that severely reduces genetic diversity (Le Corre & Kremer 1998; Clegg *et al*. 2002). Naturally recolonised populations can thus be more vulnerable to the accumulation of genetic load than reintroduced populations in species or environments with limited dispersal possibilities.

Recent years have seen increased accessibility of genomic data that provide greater power to study population structure, genetic diversity, and inbreeding (Supple & Shapiro 2018), which are important for understanding the genetic outcomes of reintroductions (Hicks *et al*. 2007; Taylor & Jamieson 2008; Wright *et al*. 2014). One such advantage of genomic data is its utility for quantifying inbreeding using runs of homozygosity (RoH). These RoH occur when breeding between individuals that share common ancestors results in offspring with stretches of homozygosity along segments of their homologous chromosomes that both parents inherited from a common ancestor (Kardos *et al*. 2015, 2016). The ability to quantify the length of RoH segments enables us to distinguish between inbreeding due to recent or more distant shared ancestors of the parents based on the distribution of RoH lengths, giving insights into the demographic history of populations (Druet & Gautier 2017; Kardos *et al*. 2017; Brüniche-Olsen *et al*. 2018).

While the success of reintroductions has been studied across a variety of taxa including fish (Drauch & Rhodes 2007), birds (Brekke *et al*. 2011), insects (White *et al*. 2017), and other ungulates (Grossen *et al*. 2018), few studies have been able to evaluate and compare their genetic consequences with those from natural population reestablishments. The wild, endemic Svalbard reindeer (*Rangifer tarandus platyrhynchus* Vrolik, 1829) subspecies, with its strong metapopulation structure (Peeters *et al*. 2020), is a biological system well suited for comparing the genetic consequences of reintroductions to those of natural recolonisations. The number of Svalbard reindeer declined drastically due to overharvesting until 1925, when they were protected, and the subspecies was extirpated from much of the Svalbard archipelago, with evidence of reindeer surviving in four isolated populations totalling ∼1000 individuals (Lønø 1959; Le Moullec *et al*. 2019). The subspecies has since largely recovered, with natural recolonisation and anthropogenic reintroductions restoring most of its former range. Accordingly, Svalbard reindeer are now abundant (∼22,000 individuals) (Le Moullec *et al*. 2019) with most populations relatively stable or increasing in size (Hansen *et al*. 2019b), and populations previously extirpated are still recovering in number (Le Moullec *et al*. 2019). As environmental conditions are rapidly changing due to climate change (Isaksen et al. 2022), including sea-ice coverage (Peeters *et al*. 2020), genetic diversity and differentiation of reindeer populations may be important for their capacity to adapt to these conditions and influence their future population dynamics. Therefore, knowledge of the genetic consequences of the Svalbard reindeer reintroduction and recolonisation events is important for understanding how the metapopulation might respond to environmental change in the future, and to contribute to a broader understanding of the genetic consequences of species reintroductions.

Here, we use whole-genome sequencing data to investigate the genetic consequences of two Svalbard reindeer reintroductions (each founded by 12 individuals (Gjertz 1995; Aanes *et al*. 2000)) and compare these to natural recolonisation processes in adjacent, comparable habitats with similar ecological conditions. Specifically, we quantify the degree to which the genetic diversity of the source population was retained through the single founder event bottleneck and subsequent rapid population growth (Kohler & Aanes 2004) associated with anthropogenic reintroduction, and whether a signature of this reintroduction could be detected in the form of longer RoH. Additionally, we investigated whether naturally recolonised populations that were not admixed would show different patterns of genetic diversity and inbreeding coefficients compared to reintroduced populations due to the compounding effects of sequential founding events during natural recolonisation.

## Results

### Sequencing

Whole-genome sequencing resulted in a mean nuclear genome sequencing depth of 3.3x for the 90 samples sequenced to a lower target coverage and 23.4x for 10 deep-sequenced samples, after all filtering (see Fig S1 for distribution of sequencing coverage). Four samples had <0.1x coverage and thus were used only for the site frequency spectrum (SFS) estimates. Genotype likelihoods for 8,255,693 variable sites were calculated from the caribou nuclear genome (Taylor et al. 2019) mapped sequence data after quality filtering. 6,309,215 of these sites remained after removing scaffolds mapping to the bovine X chromosome. Mean sequencing depth of the mitochondrial genome was >1000x.

### Admixture and principal component analyses

Admixture and principal component analysis identified clear genetic structure in the Svalbard reindeer metapopulation (Fig 1 and 2). Principal component analysis of the whole Svalbard metapopulation suggested tight clustering among a “Central Svalbard” group of populations consisting of the reintroduced (Reintroduction 1 and 2), the reintroduction source (ADV), and the naturally recolonised southern Svalbard (STH) populations (Fig 1). The major axis of variation in the PCA (PC1, Fig 1) was driven by variation between this Central Svalbard group and Mitrahalvøya (MTR), while the secondary PC axis was driven by variation between the Eastern (EST) population and both MTR and the Central Svalbard group populations, indicating strong genetic differentiation between these three groups (Fig 1). Admixture analysis further supported these patterns (Fig 2). The optimal number of genetic populations identified using the delta K method was *K*=2 for both the admixture analysis on the whole dataset and the analysis limited to the subset of populations assigned to the Central Svalbard genetic group in the Svalbard-wide *K*=2 model (Fig S2). Strongly correlated residuals between individuals in the *K*=2 models examined by EvalAdmix (Fig S3) indicate these models were a poor fit to the data and may fail to capture finer scale structure. Due to this we also present higher *K*-value models to examine finer-scale genetic structure. On the broadest scale, the *K*=2 model indicated the Central Svalbard group from the PCA share a common ancestral population distinct from the naturally recolonised MTR population, close to the reintroduction 1 site (Fig 2). The *K*=3-10 and *K*=5-10 models suggested that the remnant populations in EST and northeastern Svalbard (NE), respectively, originate from distinct ancestral populations (Fig S4). This was supported by highly correlated residuals within these populations in the models that assigned them as admixed populations (Fig S4). Individuals in the naturally recolonised Wijdefjorden (WDF) population were assigned admixed ancestry from the MTR and NE ancestral populations (models *K*=5-8), or a distinct ancestral population with some admixture with the ancestral NE population (models *K*=8-10).

**Figure 1.**
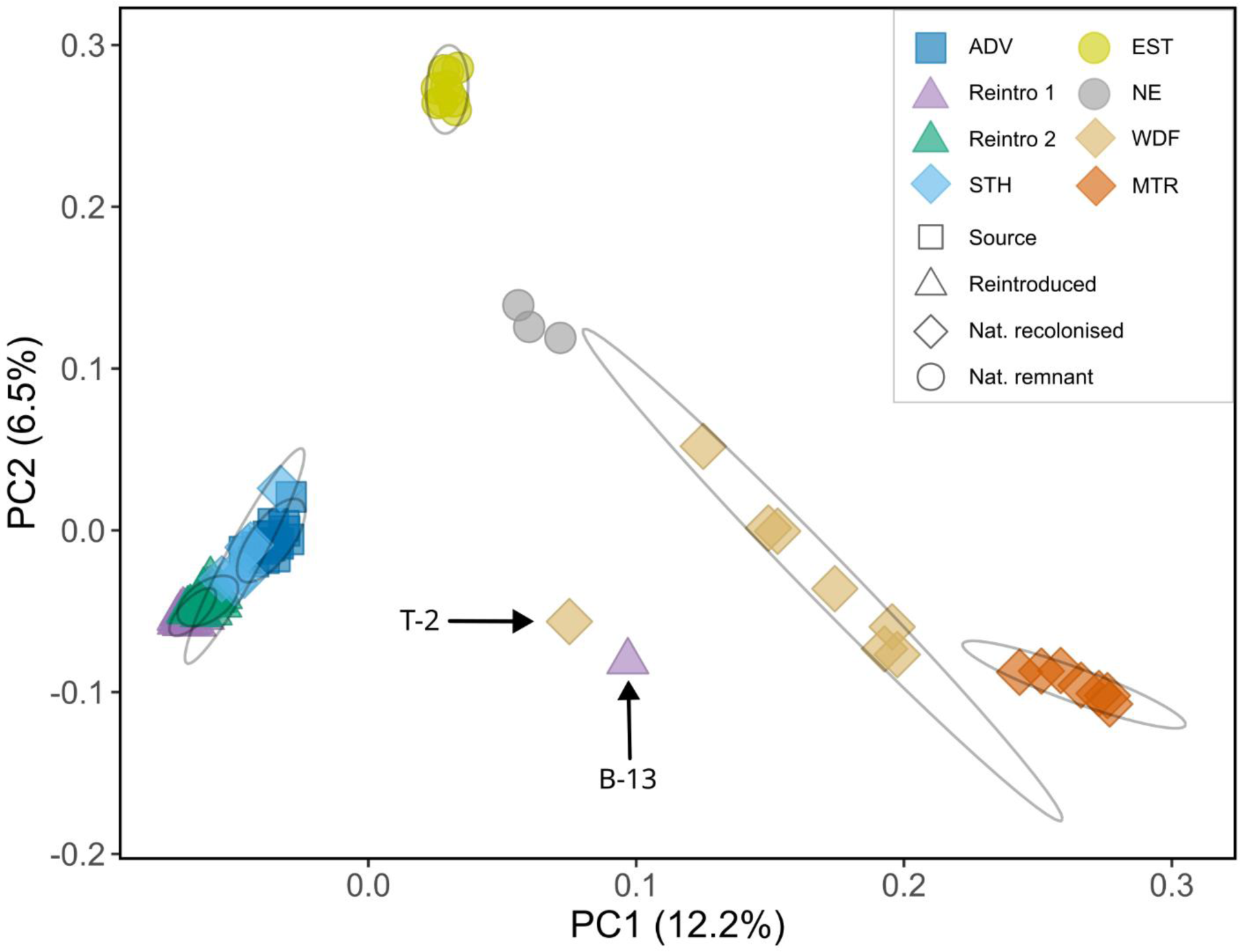
Principal component analysis plot showing PC 1 and 2. Shapes indicate type of Svalbard reindeer population and colours indicate sample population. Ellipses represent the 95% CI of the mean PC coordinates for each natural population (except NE due to too few samples) and each reintroduction group. Based on NGSadmix *K*=2 model results, two individuals (T-2 and B-13) that represented admixture between strongly differentiated populations were not included in population ellipse calculations. ADV: Adventdalen; BGR: Brøggerhalvøya; SAR: Sarsøyra; KAF: Kaffiøyra; PKF: Prins Karls Forland; DAU: Daudmannsøyra; NIF: North Isfjorden; WDF: Wijdefjorden; MTR: Mitrahalvøya; STH: Southern Spitsbergen; EST: Eastern Svalbard; NE: North East Land.

**Figure 2.**
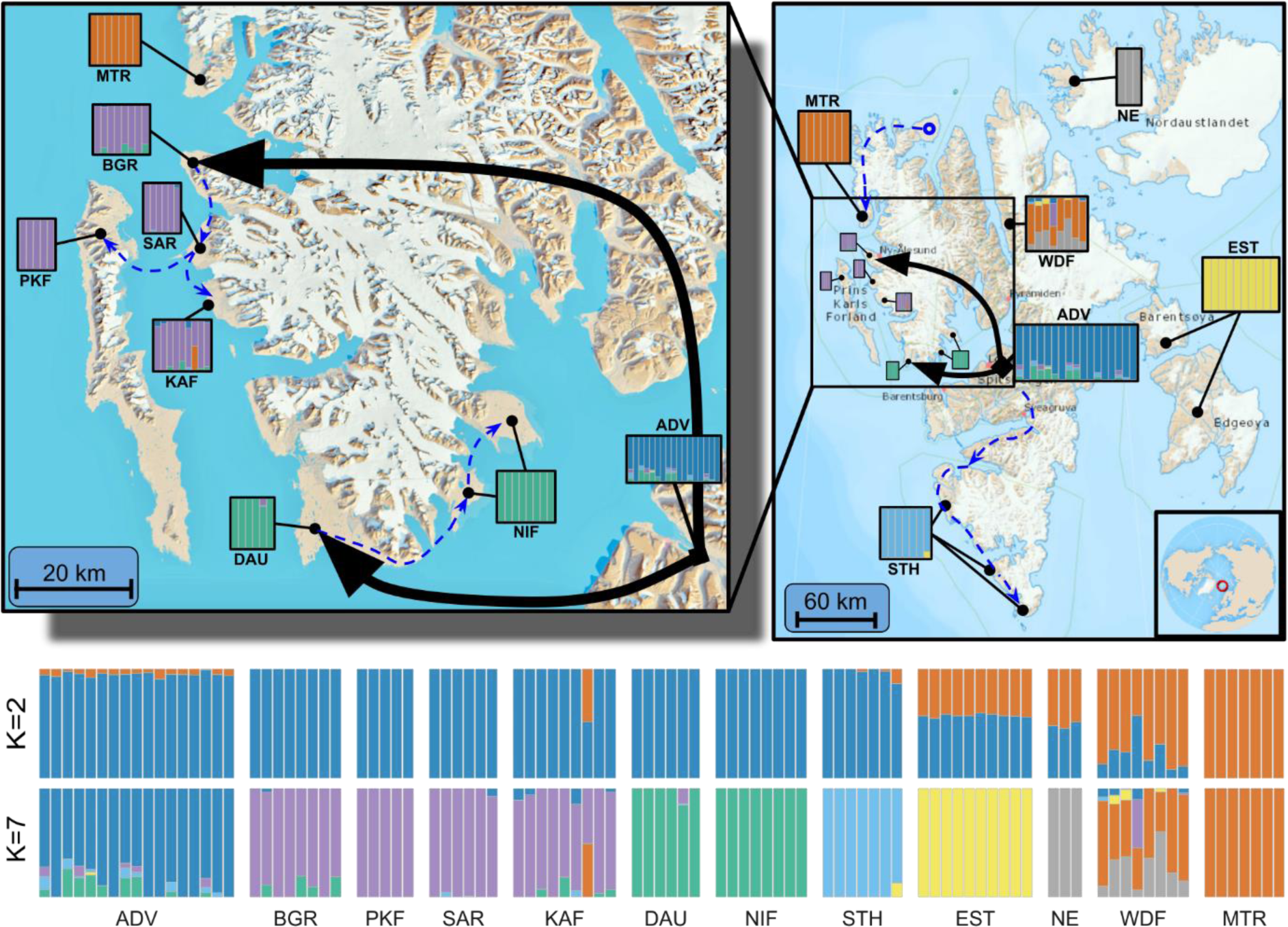
Admixture analysis results from NGSadmix analysis of Svalbard reindeer nuclear genomes. Upper: Admixture proportions for model *K*=7 (bars) and mtDNA haplogroup composition (pie charts), shown at population locations. Arrows indicate translocations for reintroduction 1 and 2; Lower: Admixture proportions for *K*=2 and *K*=7 models. Vertical bars represent individual reindeer and colours correspond to genetic cluster assignment. Black arrows indicate reintroduction translocations and dashed blue lines indicate assumed natural recolonisation routes. Maps obtained from the Norwegian Polar Institute (toposvalbard.npolar.no). ADV: Adventdalen; BGR: Brøggerhalvøya; SAR: Sarsøyra; KAF: Kaffiøyra; PKF: Prins Karls Forland; DAU: Daudmannsøyra; NIF: North Isfjorden; WDF: Wijdefjorden; MTR: Mitrahalvøya; STH: Southern Spitsbergen; EST: Eastern Svalbard; NE: North East Land.

On a finer scale, both the Svalbard-wide PCA (PC3 and PC4, Fig S5) and admixture analysis (model *K*=7, Fig 2) suggested fine-scale structure among the Central Svalbard group, confirmed by PCA and admixture analyses including only Reintroduction 1 & 2, ADV, and STH. For these fine-scale analyses, PC axes 1 and 2 (Fig S6) showed clear segregation between Reintroduction 1, Reintroduction 2, ADV, and STH. Admixture analyses showed low levels of admixture between the distinct ancestral populations corresponding to the two reintroductions, the ADV source population and the naturally recolonised STH population (*K*=4 admixture model, Fig S7). Evidence of admixture between reintroduced and natural populations was found in only one individual “B-13” from KAF (Reintroduction 1) that carried approximately 50% MTR ancestry, and one individual in WDF “T-2” that was assigned approximately 50% Reintroduction 1 and ADV ancestry. This admixture assignment was consistent with the PCA and all *K*-value admixture models (Figs 2,3 and S4).

**Figure 3.**
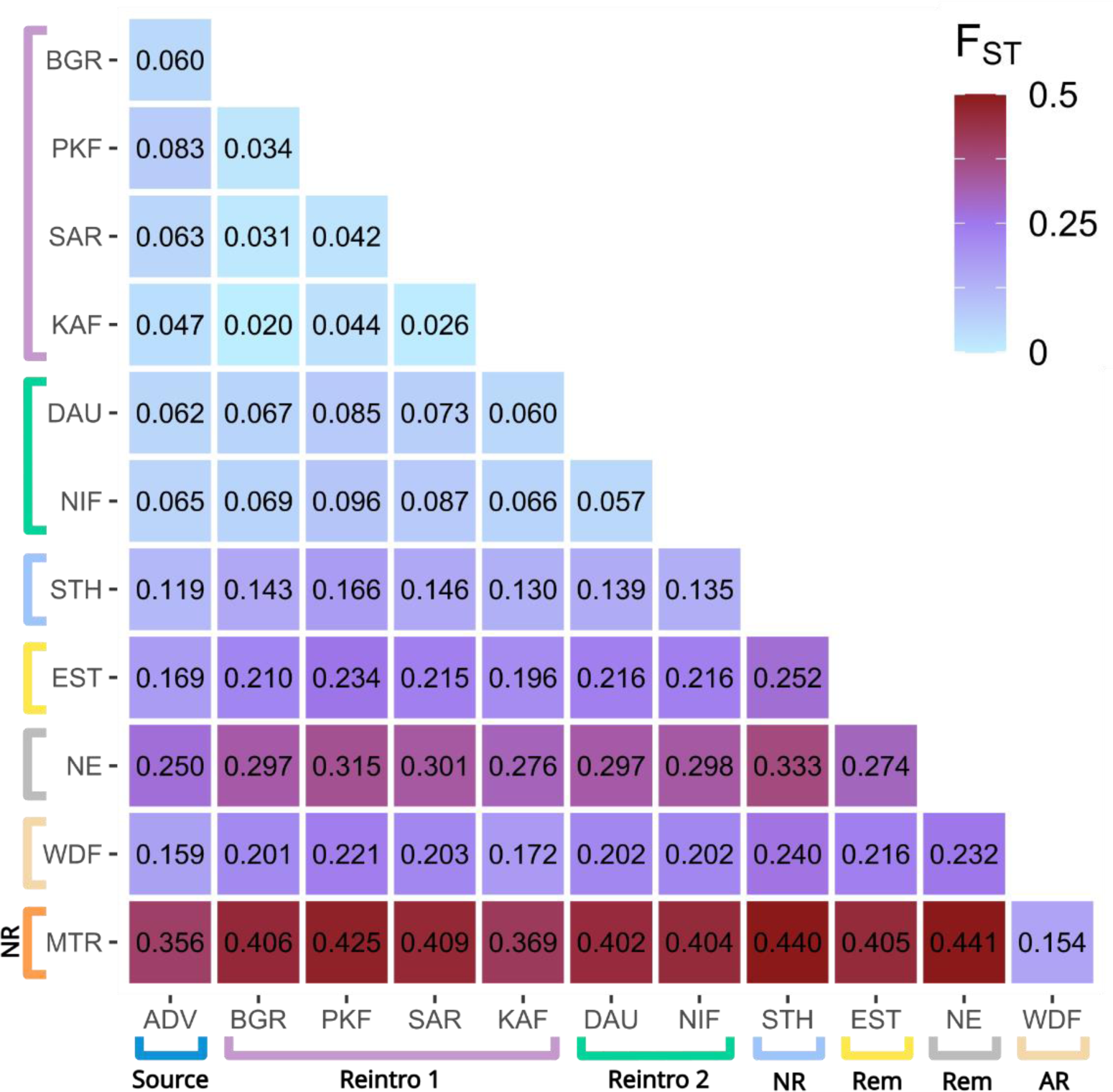
Pairwise F_ST_ heatmap for each Svalbard reindeer population based on folded SFS. Coloured brackets correspond to admixture groups assigned by the *K*=7 model. NR - Non-admixed naturally recolonised population; NR: Non-admixed naturally recolonised population; AR: Admixed naturally recolonised population; Rem: Natural remnant population. ADV: Adventdalen; BGR: Brøggerhalvøya; SAR: Sarsøyra; KAF: Kaffiøyra; PKF: Prins Karls Forland; DAU: Daudmannsøyra; NIF: North Isfjorden; WDF: Wijdefjorden; MTR: Mitrahalvøya; STH: Southern Spitsbergen; EST: Eastern Svalbard; NE: North East Land.

### F_ST_ analysis

Pairwise F_ST_ estimates showed very strong genetic structure, and largely supported admixture and PCA results (Fig 3). Populations assigned in admixture analyses to the same reintroduction showed lower pairwise F_ST_ values between each other than populations assigned to the other reintroduction. Similar levels of genetic differentiation were found in comparisons between the source population and both groups of reintroduced populations, and between the two groups of reintroduced populations. The naturally recolonised population at MTR was clearly the most genetically distinct, showing extremely high differentiation (>0.35) to all other populations except Widefjorden.

### Mitochondrial haplotype diversity

We detected 38 variant sites among the 96 full mtDNA genomes, comprising 16 unique haplotypes (Table 1, Fig 4). Haplotypes could be grouped into seven distinct haplogroups with a maximum of two substitutions separating each haplotype from its nearest neighbour **(Fig 4C)**. Mitochondrial DNA haplotype diversity showed a similar pattern to nuclear genetic analyses. Haplotypes in MTR, NE, and EST were not found in any other population except WDF, which carried a mixture of haplotypes found in every population except STH and NE, plus one highly unique haplotype **(Fig 4A)**. Overall, populations with Central Svalbard ancestry (ADV, STH, and reintroductions) shared similar haplogroups, with the notable exception that the most common haplogroup among reintroduced populations and STH was not found in the ADV source population but instead in EST. Within this shared haplogroup, the haplotypes found in STH and the reintroduced populations were mutually exclusive to those in EST, but differed by as little as a single mutation **(Fig 4B)**. The populations KAF and NIF, from Reintroduction 1 and 2 respectively, carried haplotypes belonging to haplogroups not found in other reintroduced populations (Fig 4a). The KAF individual admixed with MTR (Fig 2) carried a unique haplotype from the MTR haplogroup, and almost half of the NIF samples carried haplotypes in a haplogroup otherwise found only in ADV (Fig 4A).

**Figure 4.**
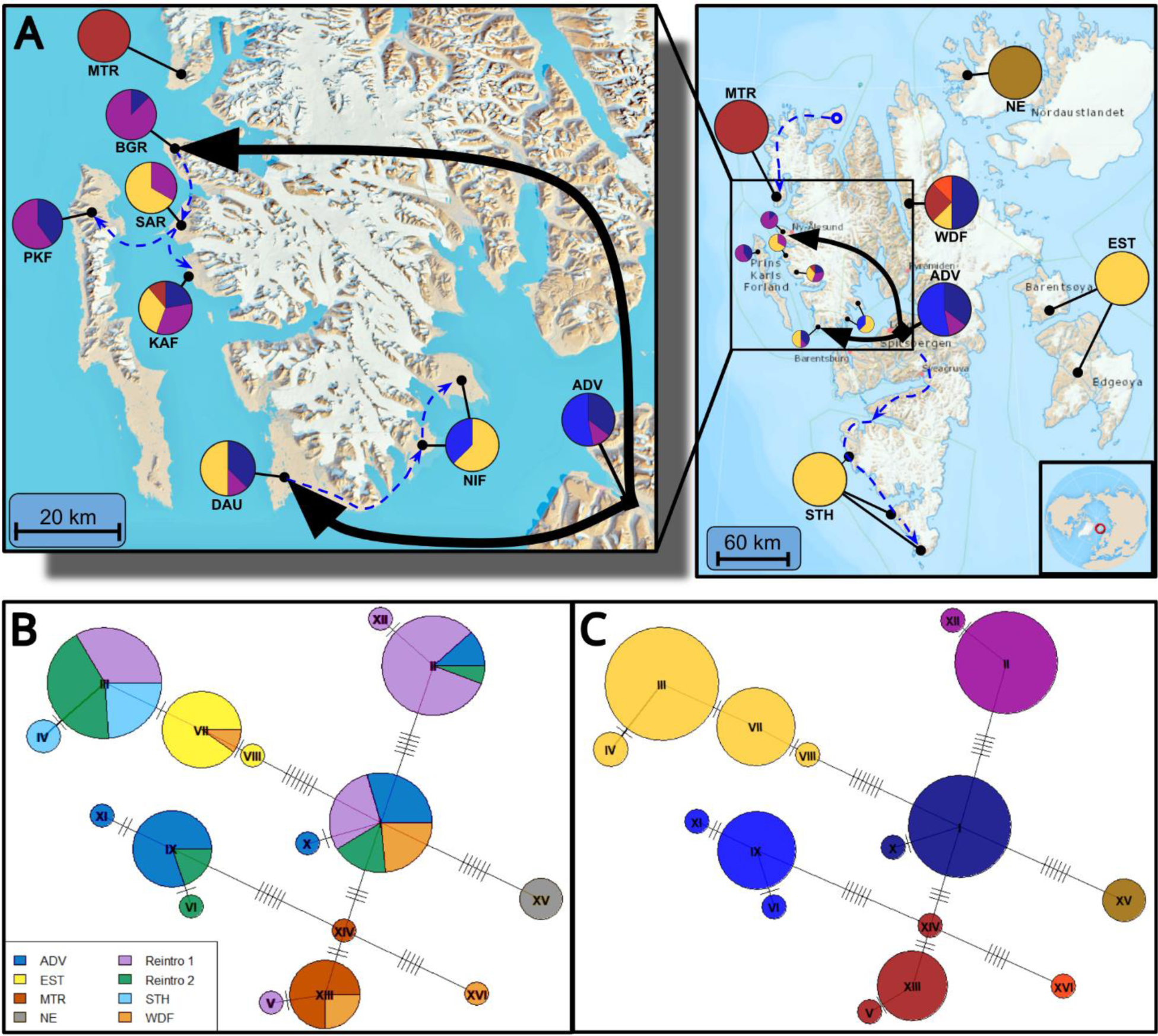
Svalbard reindeer mtDNA haplotype analysis. A) Population haplogroup composition. Colours indicate haplogroups shown in panel C. Black arrows indicate reintroduction translocations and dashed blue lines indicate assumed natural recolonisation routes; B) Median joining mitochondrial haplotype network constructed using uncorrected number of nucleotide differences (colours represent populations); C) Grouping of haplotypes into haplogroups connected by links with less than 3 nucleotide differences. Maps obtained from the Norwegian Polar Institute (toposvalbard.npolar.no).ADV: Adventdalen; BGR: Brøggerhalvøya; SAR: Sarsøyra; KAF: Kaffiøyra; PKF: Prins Karls Forland; DAU: Daudmannsøyra; NIF: North Isfjorden; WDF: Wijdefjorden; MTR: Mitrahalvøya; STH: Southern Spitsbergen; EST: Eastern Svalbard; NE: North East Land.

Haplotype richness was strongly correlated to mean population genome-wide heterozygosity (pearson correlation r=0.86, p=0.013). Each of the two reintroduction groups (populations combined) had similar haplotype richness to ADV, and higher than all other natural populations except WDF (Table S2 and Fig S8).

### Heterozygosity analysis

The median genome-wide heterozygosity for all individuals in this study with coverage >2.5x was 3.02×10^-4^. Populations originating from the first and second reintroductions had slightly lower median genome-wide heterozygosities (3.13×10^-4^, IQR 4.59×10^-5^ and 3.19×10^-4^, IQR 4.94×10^-5^ respectively) than the ADV source population (genome-wide heterozygosity 3.31×10^-4^, IQR 1.68×10^-5^, Fig 5), but overall, there was weak evidence that heterozygosity was different between reintroduced individuals (both populations combined) and those from ADV (Mann-Whitney U test, W=483, P=0.081). In contrast to the remnant natural and admixed recolonised populations with intermediate heterozygosities (EST, NE, and WDF), the two non-admixed naturally recolonised populations MTR (median 1.89×10^-4^, IQR 7.14×10^-5^) and STH (median 2.34×10^-4^, IQR 6.37×10^-5^) had very low heterozygosity. These recolonised populations had markedly lower than the reintroduced populations combined (W=240, P<0.001 and W=235, P<0.001 respectively).

**Figure 5.**
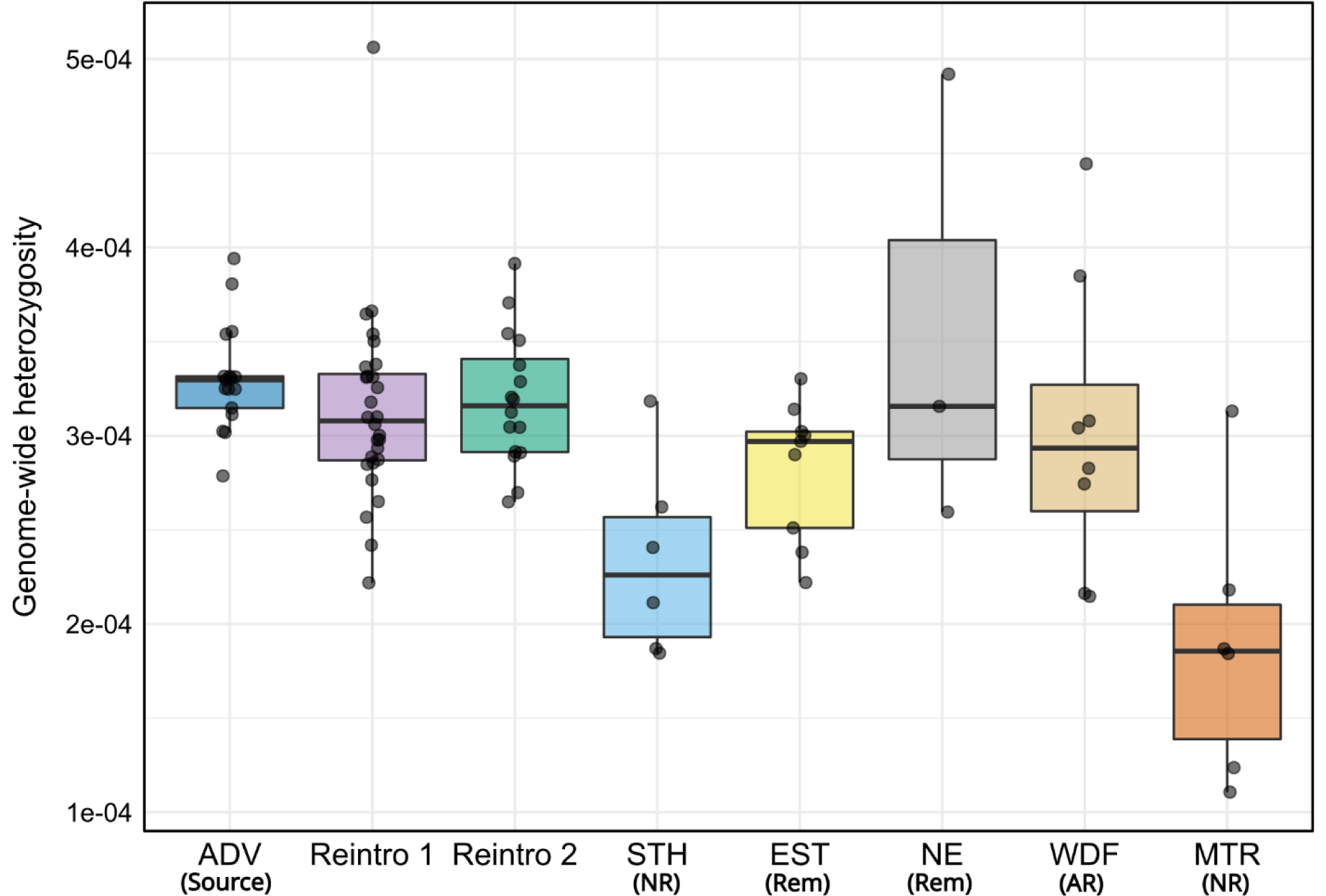
Svalbard reindeer genome-wide heterozygosity estimates using sequence data downsampled to 2.5x coverage. Reintroduced populations grouped into Reintroduction 1 and 2 based on the admixture analyses. NR: Non-admixed naturally recolonised population; AR: Admixed naturally recolonised population; Rem: Natural remnant population. ADV: Adventdalen; BGR: Brøggerhalvøya; SAR: Sarsøyra; KAF: Kaffiøyra; PKF: Prins Karls Forland; DAU: Daudmannsøyra; NIF: North Isfjorden; WDF: Wijdefjorden; MTR: Mitrahalvøya; STH: Southern Spitsbergen; EST: Eastern Svalbard; NE: North East Land.

### Inbreeding

We detected 21,990 RoH longer than 0.5 Mbp across the 83 genomes included in the analysis, ranging from 0.5 Mbp to 12.76 Mbp in length and covering between 3% to 59% of individual’s genomes (Fig 6). Both reintroduced populations showed a higher mean inbreeding coefficient (Reintroduction 1 F_ROH_ 0.236 ± 0.063 SD, Reintroduction 2 F_ROH_ 0.237 ± 0.033) relative to their source population in Adventdalen (F_ROH_ 0.186 ± 0.042). The higher inbreeding in reintroduced populations was due to greater coverage of moderate length RoH (between 1 - 8 Mbp) that resulted in increased median RoH lengths (Fig 6), indicating a reduced effective population size in the more recent past compared to the ADV source population.

**Figure 6.**
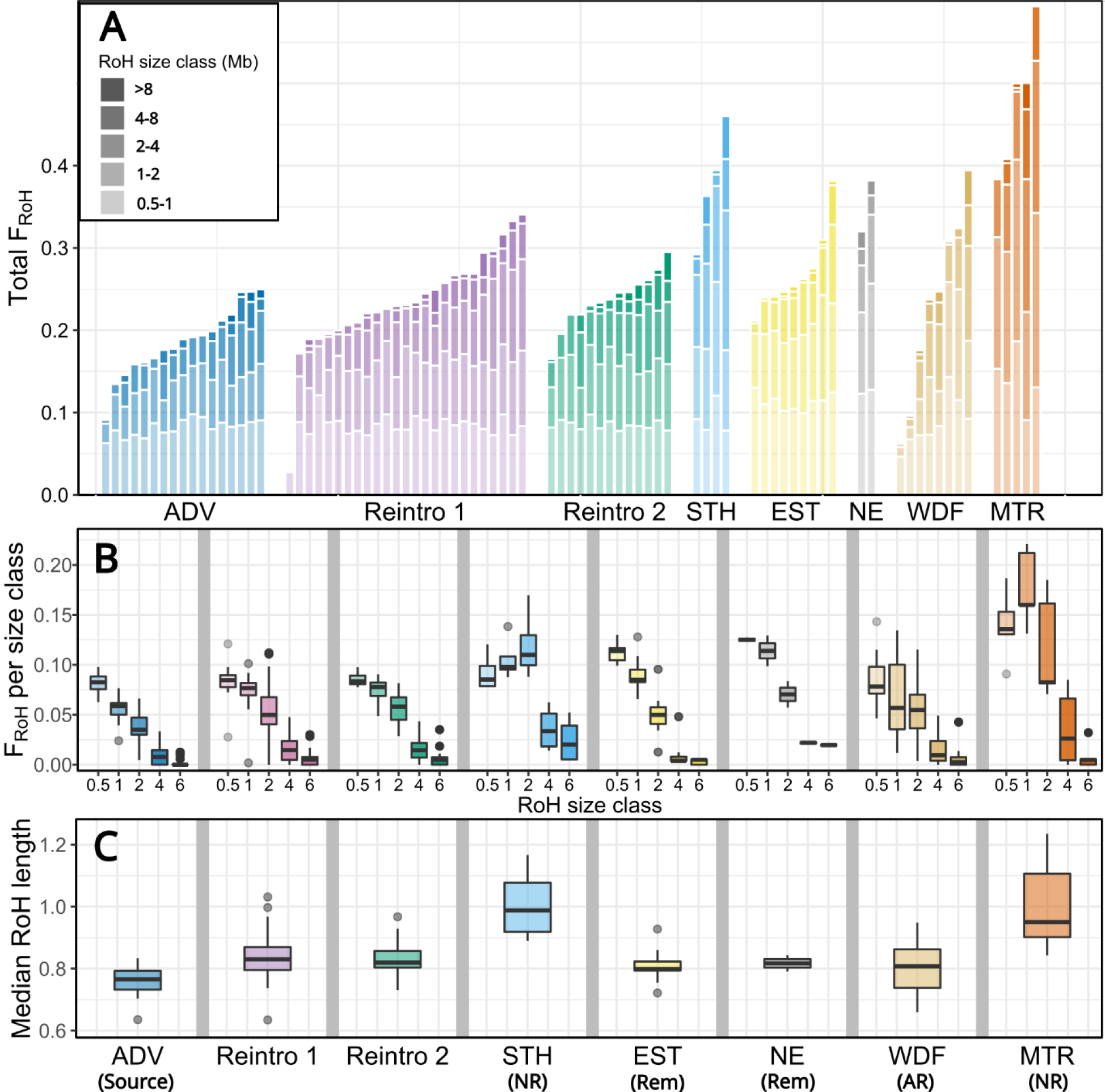
A) Cumulative total F_ROH_ from the five Runs of Homozygosity (RoH) size classes (0.5-1 Mbp, 1-2 Mbp, 2-4 Mbp, 4-8 Mbp, and >8 Mbp) with each bar representing an individual Svalbard reindeer genome; B) Proportion of individual genomes within RoH of each size classes; C) Median RoH lengths of individuals in each population. NR: Non-admixed naturally recolonised population; AR: Admixed naturally recolonised population; Rem: Natural remnant population. ADV: Adventdalen; BGR: Brøggerhalvøya; SAR: Sarsøyra; KAF: Kaffiøyra; PKF: Prins Karls Forland; DAU: Daudmannsøyra; NIF: North Isfjorden; WDF: Wijdefjorden; MTR: Mitrahalvøya; STH: Southern Spitsbergen; EST: Eastern Svalbard; NE: North East Land.

Non-admixed naturally recolonised populations MTR and STH had the highest inbreeding coefficients (mean F_ROH_ 0.477 ± 0.084 and 0.377 ± 0.07, respectively) and longest median RoH lengths (Fig 6). We found the strongest signals of recent inbreeding in STH, with a high proportion of individuals’ genomes covered by RoH >4 Mbp relative to other populations and a peak in F_ROH_ in the 2-4 Mbp class (Fig. 6B). In MTR, F_ROH_ was highest in the 1-2 Mbp class with a large contribution from all other size classes <8 Mbp relative to other populations, but very long (>8 Mbp) RoH were rare. The admixed naturally recolonised population in WDF had the highest variation in total F_ROH_ consistent with high variation in the proportion of individuals’ shared ancestry. The two remnant natural populations EST and NE both had high F_ROH_ in short RoH classes compared to the ADV population, with EST having low F_ROH_ in intermediate and long size classes comparable to ADV, and the two samples from NE showing relatively high F_ROH_ in all size classes.

## Discussion

By analysing whole genome sequences from more than 100 Svalbard reindeer across their range, we have quantified important genomic consequences and contrasts of population recovery through anthropogenic reintroductions versus natural recolonisations in the previously severely overharvested subspecies. We found strong archipelago-wide genetic structure, including two distinct genetic clusters corresponding to two reintroductions from a common source, with little evidence for extensive admixture between reintroduced and sampled natural populations (Figs 1 - 4). Our results show that reintroduced populations also maintained comparative levels of genetic diversity to their source population (Fig 5), although bottleneck effects resulted in a small increase in the length and coverage of RoH across the genomes of reintroduced individuals (Fig 6). This contrasted strongly to non-admixed naturally recolonised populations, which had markedly lower genetic diversity and greater proportion of their genomes comprising RoH. This suggests that non-admixed, naturally recolonised populations may be more vulnerable to the accumulation of genetic load and loss of adaptive variation than reintroduced populations, even when the latter originate from just a handful of individuals.

### Effect of reintroduction versus recolonisation on genome-wide diversity

We estimated genome-wide heterozygosity and analysed the distribution of RoH lengths to separate the contribution of ancient and more recent demographic history, associated with reintroduction and recolonisation, to patterns of genome-wide variation. Our analyses identified a weak signal of a population bottleneck in both reintroductions, which had higher total inbreeding coefficients and longer RoH, but no significant reduction in average heterozygosity compared to the source population. Similar genomic signatures of reintroductions have previously been identified, including in European bison *Bison bonasus* (Druet *et al*. 2020), Magpie-robins *Copsychus sechellarum* (Cavill *et al*. 2022), and ibex *Capra ibex* (Grossen *et al*. 2018). The increased inbreeding in reintroduced populations was attributable to a greater proportion of the genome in RoH >1 Mbp, including in the 1-2 and 2-4 Mbp range. These size classes reflect shared ancestry approximately 25-50 and 12-25 generations ago, i.e. before the time of the reintroduction translocation, given an average generation time of 5-6 years. We found only a small increase in the coverage of long RoH (4-8 Mbp), which is the expected length of RoH caused by shared ancestors 6-12 generations ago (around the time of the reintroduction), indicating our analysis may have underestimated RoH lengths. However, generation time may have been reduced in the post-reintroduction period of strong population growth due to an abundance of resources resulting in an earlier age of first reproduction and high fecundity (Giaimo & Traulsen 2022). Nevertheless, a low frequency of long RoH relative to naturally recolonised populations suggests lower levels of recent inbreeding among reintroduced individuals. The founding population sizes of twelve Svalbard reindeer have thus been sufficient to maintain most of the heterozygosity of the source population and avoid serious accumulation of inbreeding in both reintroductions, both of which are a key concern for reintroduced populations (Weeks *et al*. 2011; Frankham *et al*. 2017).

We found fewer long RoH in reintroduced Svalbard reindeer than reported for alpine ibex (Grossen *et al*. 2018) and European bison (Druet *et al*. 2020), and our results indicate that, like the source population (ADV), short RoH (i.e. inbreeding due to ancient demography) still contribute the most to the total inbreeding coefficients of reintroduced individuals. Rapid population growth immediately after reintroduction (Aanes *et al*. 2000) and the extensive overlap between generations in reindeer are both characteristics that reduce loss of genetic diversity after population bottlenecks (Nei *et al*. 1975; Allendorf 1986). Furthermore, past and possibly ongoing dispersal among the secondary sub-populations recolonised from the initial reintroduced populations (Hansen *et al*. 2010; Stien *et al*. 2010) may have buffered against the effects of sequential founder events. Similar outcomes have been observed in reintroductions of a range of vertebrate and invertebrate species where rapid population growth occurred after reintroduction (Hicks *et al*. 2007; Brekke *et al*. 2011; Murphy *et al*. 2015; White *et al*. 2017). In particular, populations of other ungulates colonised by only a few founding individuals have been shown to retain high levels of heterozygosity as a result of overlapping generations and rapid population expansion (Kaeuffer *et al*. 2007; Kekkonen *et al*. 2012). In contrast, populations reintroduced with few founders that remained small for several generations have shown a pronounced reduction in genetic diversity (Williams *et al*. 2002; Wisely *et al*. 2008).

In contrast to the reintroductions, the natural recolonisation of STH and MTR resulted in populations with very low nuclear genome-wide heterozygosity, mtDNA haplotype diversity, and high total inbreeding coefficients with a longer distribution of RoH. In STH, high F_RoH_ across the 1-2, 2-4, and 4-8-Mbp size classes, with a peak in the 2-4-Mbp range, indicates small population size around the same time as the reintroductions (i.e. during the recolonisation period) (Fig 6B). STH also has the highest proportion of genomes within RoH >4 Mbp, indicating bottlenecks or small population size more recently than in the reintroduced populations. Long RoH may be particularly significant to conservation as they represent young haplotypes that have been exposed to less selection than older haplotypes, making them more likely to harbour deleterious mutations (Bortoluzzi *et al*. 2020) and have a greater impact on fitness (Szpiech *et al*. 2013; Stoffel *et al*. 2021). The very high frequency of short RoH in the MTR genomes suggests this population originates from a source with a historically small population size (i.e. prior to recolonisation), and high coverage of moderate length RoH indicates bottlenecks or small population size during the last 25 generations (i.e. during recolonisation). Both ADV, the two reintroduced populations, and the naturally recolonised STH had similar average proportion of genomes within RoH <1 Mbp, indicating similar demographic histories >50 generations ago, consistent with these populations sharing a common origin as indicated by our admixture results. Therefore, differences in patterns of genome-wide diversity in STH and the reintroduced populations likely reflect differences between anthropogenic reintroductions and natural recolonisation, rather than differences in genetic diversity between their ancestral populations. The extremely low levels of heterozygosity and high inbreeding levels in MTR are thus likely a result of a source population with historically small population size that harboured little genetic diversity prior to recolonisation, in addition to more recent population bottlenecks associated with the recolonisation process. Indeed, this is consistent with the reported population size of reindeer in the isolated North-West Spitsbergen area being only 2-300 individuals, 30 years after the historical overharvesting was ended (Lønø 1959).

The decreased heterozygosity and increased frequency of longer RoH in naturally recolonised populations likely reflect multiple bottlenecks from a sequential recolonisation process (Peeters *et al*. 2020) (potentially involving few dispersing individuals) that was avoided by the direct anthropogenic reintroduction. Sequential dispersal and establishment in isolated peninsulas and valleys during the recolonisation of STH and MTR may have caused cumulative effects from multiple founder-events, reducing effective population sizes and eroding genetic diversity (Le Corre & Kremer 1998; Clegg *et al*. 2002; Pruett & Winker 2005), resulting in increased inbreeding and long RoH (Grossen *et al*. 2018).

Our inferences regarding the timing of past demography based on RoH length distributions should be considered only as relative between populations and may not accurately reflect the demographic history in absolute number of generations. Inferences of demographic history using RoH length distributions are imprecise because spatial and temporal variation in generation times and the random nature of recombination result in high variation around the mean expected length of RoH (Druet & Gautier 2017). Moreover, sequencing coverage, sequencing error rates, biased genotype likelihood estimates as well as filtering, and parameter settings can all affect estimates of heterozygosity (Fuentes-Pardo & Ruzzante 2017; de Jager *et al*. 2021; Sánchez-Barreiro *et al*. 2021), and thus RoH length and frequency (Duntsch *et al*. 2021). We down-sampled sequence data to allow unbiased comparison between samples and populations with varying levels of coverage, and to maximise the sample size for both the genome-wide heterozygosity and RoH analyses. This may limit the direct comparison of our estimates of heterozygosity to those from other studies using higher-coverage sequence data.

RoH analysis using low coverage sequence data is also sensitive to filtering, and may falsely identify multiple short RoH as a single longer RoH, or miss RoH altogether (Duntsch *et al*. 2021). However, the shorter than expected distribution of RoH in reintroduced individuals, given the known timing of the reintroduction bottleneck, suggests our analysis underestimated RoH lengths, and this is consistent with an excess of short (<50-Kbp) gaps between RoH in our data, breaking up otherwise long RoH (Fig S9). Several factors could break up true RoH and contribute to downwardly biased RoH length distributions: (1) Sequencing errors resulting in false heterozygosity can break up RoH, and will have a larger effect on accurate identification of longer RoH (MacLeod *et al*. 2013); (2) Any RoH spanning the genome assembly scaffold edges will be broken up (Brüniche-Olsen *et al*. 2018), thus lacking a chromosome-level genome assembly, we only included scaffolds longer than 10 Mbp in RoH analyses; (3) Errors in mapping sequence reads or structural variation between the caribou reference genome assembly and the Svalbard reindeer genome could also break up long RoH. Reanalysing RoH after allowing one <50-Kbp gap within each 1-Mbp segment (similar to Wilder *et al*. (2022)) gave qualitatively similar results but showed more long RoH which aligned more closely with the known timing of the reintroduction bottleneck (Fig S10).

### Genetic structure within the Svalbard reindeer metapopulation

Admixture, F_ST_, and mitochondrial haplotype analyses identified strong genetic structure across the archipelago, in some cases even over short geographical distances, confirming patterns identified with microsatellite data in Peeters *et al*. (2020). Such genetic structure is typical of ungulate populations with a history of population fragmentation and bottlenecks due to past harvesting pressure (Williams *et al*. 2002; Haanes *et al*. 2010). On a finer scale, this study reveals population structure within the Central Svalbard group, i.e. between the source and reintroduced populations, and among reintroduced populations. The two distinct genetic clusters among reintroduced populations corresponded to the two separate reintroductions to isolated peninsulas on the west coast of Spitsbergen. Pairwise F_ST_ estimates reveal both reintroductions have resulted in a similar degree of genetic divergence from the source population. Founder effects and subsequent genetic drift commonly induce structure between reintroduced populations and their sources, typically reflecting isolation from the source population and subsequent genetic drift (Williams *et al*. 2002; Latch & Rhodes 2005; Brekke *et al*. 2011; Andersen *et al*. 2014; Grossen *et al*. 2018).

Close genetic clustering of multiple subpopulations colonised from a common reintroduced founder population is characteristic of populations manipulated by reintroduction programmes (Andersen *et al*. 2014; Grossen *et al*. 2018). We found only weak genetic structure among populations originating from the first reintroduction, except for the rather isolated island PKF, which population showed little admixture with other reintroduced populations (Fig S7), reflecting low dispersal across the sea (Peeters *et al*. 2020). Population monitoring after the reintroduction to BGR (Reintroduction 1) recorded substantial movement between BGR, SAR, and KAF (Hansen *et al*. 2010; Stien *et al*. 2010), but GPS collar data suggest that such exchange of individuals is now rare (Pedersen, Hansen and Le Moullec, unpubl. data). This is consistent with the observed lack of fjord ice in recent decades.

Our results indicate little gene flow between reintroduced populations and sampled natural populations. Only one individual from a reintroduced population was identified as admixed with a natural population, with admixture proportions consistent with an F1 offspring resulting from a mating between individuals in reintroduction 1 and MTR genetic clusters. This individual carried a unique haplotype that differs by only a single mutation from MTR haplotypes, suggesting female dispersal from the north. MTR and BGR, the closest population sampled in reintroduction 1, are separated by only 15 km across the mouths of Kongsfjorden and Krossfjorden, a span of water which has rarely or never frozen over since the reintroduction (Pavlova *et al*. 2019; Urbański & Litwicka 2022). This lack of sea ice as a movement corridor, in combination with tide-water glaciers and steep mountains inhibiting alternative dispersal routes, has likely prevented gene flow and contributed to the extreme degree of genetic differentiation between these geographically proximate populations. On the other hand, it is likely that these populations were more closely related in the past, i.e. before the local extirpations due to overharvest and the subsequent reintroduction from central Spitsbergen (to BGR) and recolonisation from the North (to MTR). This illustrates how both reintroductions and recolonisation may cause dramatic changes in population-genetic structuring and diversity.

An exception of the clearly separated reintroduced versus naturally recolonised populations occurred along the northern side of Isfjorden in central Svalbard (NIF). We found stronger genetic structure among the two sampled populations in the second reintroduction group (DAU and NIF). Our mtDNA analyses also suggest that introgression possibly occurred from a westward natural recolonisation from an unsampled population carrying a haplotype not sampled in other populations, possibly facilitated by more frequent sea ice in the inner parts of the fjord (Muckenhuber *et al*. 2016). mtDNA haplotypes found in both reintroduction groups and STH were absent from our ADV source population samples, which could also be consistent with introgression from an unsampled natural population. However, these haplotypes may have been present in ADV at the time of reintroduction, but if so it is unclear whether they existed only at low frequencies and increased in reintroduced populations due to founder effects, or if there has been a significant change in the mtDNA haplotype diversity in the ADV source population.

### Implications for conservation and management

Small or bottlenecked populations are at risk of reduced fitness due to the accumulation of genetic load (i.e. increased frequency and fixation of recessive deleterious mutations), making it an important consideration in conservation biology (van Oosterhout 2020; Kardos *et al*. 2021; Bertorelle *et al*. 2022). Additionally, severe or extended bottlenecks are expected to reduce genome-wide diversity, including functional genetic variation potentially important for the long-term adaptive potential of populations (Frankham 2005; Kardos *et al*. 2021). Severely bottlenecked recolonised or reintroduced populations of ibex have shown increased realised genetic load compared to less severely bottlenecked populations (Grossen *et al*. 2020). Similarly, bottlenecked populations of corvids *Corvus* spp (Kutschera *et al*. 2020), Montezuma quail *Cyrtonyx montezumae* (Mathur & DeWoody 2021), and rattlesnakes *Sistrurus* spp (Ochoa & Gibbs 2021) show higher realised genetic load than larger populations. Thus, the relatively mild bottleneck effects of the Svalbard reindeer reintroductions suggest they are likely to have retained more functional variation and accumulated less realised load than the natural recolonisations, which probably experienced more severe and repeated bottlenecks. However, while some of the naturally recolonised populations may be at higher risk of reduced fitness due to increased realised genetic load, these populations may also carry fewer highly deleterious recessive mutations due to purging (Glémin 2003; Grossen *et al*. 2020), and inbreeding may have less effect on individual fitness (Mathur & DeWoody 2021).

Several management practice recommendations have been put forward to give general guidelines for the number of individuals that need to be reintroduced, and the amount of gene flow required to maintain genetic diversity. For example, approximately 20 effective founders (Willis & Willis 2010) and one effective migrant per generation (Vucetich & Waite 2000) have been viewed as sufficient in mammal populations. Despite founder population sizes lower than this, population surveys have shown that both reintroduced and naturally recolonised Svalbard reindeer populations have been expanding (Le Moullec *et al*. 2019), meaning any increase in realised genetic load is not yet severe enough to prevent continued population growth.

Svalbard reindeer face rapid environmental change (Hansen *et al*. 2019b) that may have implications for the fitness and viability of populations in future. Thus far, most populations of Svalbard reindeer have experienced a net gain from climate change effects, with a warmer and longer snow-free season leading to increased survival, reproduction and abundances (Albon *et al*. 2017; Hansen *et al*. 2019b; Loe *et al*. 2021). However, an increase in winter precipitation or “rain-on-snow” events increases ground ice cover during winter (Peeters *et al*. 2019) which limits access to winter forage (Hansen *et al*. 2010; Loe *et al*. 2016), occasionally destabilising reindeer population dynamics (Kohler & Aanes 2004; but see Stien *et al*. 2010 and Hansen *et al*. 2019a). Any short- or medium-term evolutionary responses to such environmental changes will likely depend on sufficient standing genetic variation for natural selection to act upon (Carlson *et al*. 2014). The potentially genetically depleted naturally recolonised populations may have limited ability to adapt to these changes. There is also the potential for environmental changes to accentuate the fitness effects of realised genetic load. Inbreeding depression (i.e. the fitness effects of realised genetic load) can be more severe in stressful environmental conditions (Armbruster & Reed 2005; Fox & Reed 2011). Therefore, previously benign or mildly deleterious variation in reindeer populations may have more severe fitness effects in the future. The Svalbard Archipelago is also experiencing a rapid decline in sea-ice, which acts as a movement corridor important for facilitating gene flow in Svalbard reindeer (Peeters *et al*. 2020). Future sea-ice reductions may further isolate and fragment reindeer populations, so despite the overall population expansion of Svalbard reindeer, accumulation of inbreeding and further loss of diversity in already bottlenecked populations should remain a concern.

Our results suggest that anthropogenic reintroductions can sometimes be more effective than natural recolonisations in establishing genetically healthy populations, especially under increased anthropogenic landscape fragmentation, such as that due to sea-ice reductions. This is likely due to a higher number of (potentially more genetically diverse) founders and the avoidance of sequential population bottlenecks that may be an inherent part of the natural recolonisation process in fragmented landscapes. Much of the Svalbard archipelago has been recently recolonised by reindeer (i.e, within the last century; Le Moullec *et al*. 2019), and further sampling may shed more light on whether low diversity and high inbreeding is a general pattern across recolonised populations of Svalbard reindeer, and indeed in other fragmented species. Future research may benefit from fitness and phenotypic data, modern and historical reindeer samples, and molecular methods of quantifying both functional variation and genetic load (Bertorelle *et al*. 2022) to better understand the status of Svalbard reindeer populations and, more generally, how reintroduction and natural recolonisation processes affect genetic load, inbreeding depression, and potential adaptive variation in the wild.

## Materials and Methods

### Study area

The Norwegian high-arctic Svalbard archipelago lies between the Arctic Ocean, the Barents Sea and the Greenland Sea, approximately 700 km north of mainland Norway (76-81°N, 10-35°E). Only 16% of the archipelago’s land area comprises vegetated peninsulas and valleys (Johansen *et al*. 2012), which are fragmented by tide-water glaciers and inland and mountains that cover the majority of the land area. Vegetation types on the archipelago include polar deserts, Northern Arctic tundra dominated by prostrate dwarf shrubs and cryptogams, and Middle Arctic tundra dominated by erect dwarf shrubs, forbs, and grasses (Jónsdóttir 2005).

### Study species

The Svalbard reindeer is an endemic subspecies that likely colonised the archipelago from Eurasia 6700 to 5000 years ago (Kvie *et al*. 2016). The subspecies is the dominant and only large herbivore in the terrestrial ecosystem, with little interspecific competition and almost non-existent predation pressure (but see Derocher *et al*. (2000) and Stempniewicz *et al*. (2021)). Reindeer were overharvested to near-extinction on Svalbard during the 19^th^ and early 20^th^ centuries, before coming under legal protection from hunting in 1925 (Le Moullec *et al*. 2019). By this time, it had been extirpated from much of its former natural range, and isolated remnant populations were largely confined to four regions: the northern, northeastern, and eastern extremes of the archipelago, as well as the central Spitsbergen region (Lønø 1959; Le Moullec *et al*. 2019). After coming under legal protection, the subspecies began to recover but was still absent from much of its range in the 1970s, including the west coast of Spitsbergen. In 1978, 12 surviving individuals (nine females and three males) were translocated from Adventdalen in central Spitsbergen to Brøggerhalvøya on the west coast (Fig 2, (Aanes *et al*. 2000)). In 1984-85 a second translocation reintroduced 12 individuals to Daudmannsøyra, on the north-western edge of Isfjorden (Gjertz 1995), however there is no population monitoring data to confirm these survived and established the current population. The reindeer population size at Brøggerhalvøya has been annually monitored since the reintroduction (Aanes *et al*. 2000; Hansen *et al*. 2019b). This has recorded the population’s rapid expansion after translocation (from 12 individuals in 1978 to ∼360 individuals in 1993), until a combination of high population density and poor winter conditions triggered a population crash (∼80 individuals in 1994) and migration to recolonise the nearby peninsulas of Sarsøyra (1994) and Kaffiøyra (1996) to the south, and Prins Karls Forland island (∼1994) to the west (Gjertz 1995; Aanes *et al*. 2000).

Reindeer populations have since then recolonised most of their former range naturally, including southern Spitsbergen, the north coast of Isfjorden (to the east of the reintroduced population at Daudmannsøyra), the north-west coast south to Mitrahalvøya, and Wijdefjorden in north-central Spitsbergen (Le Moullec *et al*. 2019). Populations at Mitrahalvøya, Wijdefjorden, and Southern Spitsbergen appear naturally recolonised from remnant populations, while the origins of the populations along North Isfjorden are unclear, but likely originated from the second reintroduction (Daudmannsøyra) and possibly admixed with naturally recolonising individuals (Peeters *et al*. 2020). Genetic evidence suggests the Svalbard reindeer metapopulation has low levels of genetic diversity (Kvie *et al*. 2016; Weldenegodguad *et al*. 2020) and shows strong population structure (Côté *et al*. 2002; Peeters *et al*. 2020), reflecting a history of population bottlenecks and the largely philopatric nature of the species with no large scale migration (Hansen *et al*. 2010).

### Sample collection

Genetic data were generated from tissue samples (ear, antler, bone, or fur) collected in 2014-2018 from 100 individual reindeer originating from twelve (sub)populations on the Svalbard archipelago (Fig 2, Table S3 Table S1). Based on the extirpation locations reported in Lønø (1959), we categorised these populations as either putative reintroduced or naturally recolonised extirpated populations, or remnant non-extirpated populations. These included six populations believed to have originated from the two translocations (Gjertz 1995; Aanes *et al*. 2000): (1) Brøggerhalvøya (BGR, the initial reintroduction site), Sarsøyra (SAR), Kaffiøyra (KAF), and Prins Karls Forland (PKF) from the first translocation (hereafter collectively referred to as “Reintroduction 1”), and (2) Daudmannsøyra (DAU, the second reintroduction site) and North Isfjorden (NIF) from the second translocation (hereafter referred to as “Reintroduction 2”). Samples were also collected from the source population of the reintroductions (Adventdalen, ADV), from two other remnant populations (Eastern Svalbard (EST), and North East Land (NE)), and the naturally recolonised populations (Mitrahalvøya (MTR), Southern Spitsbergen (STH), Wijdefjorden (WDF). Except those from Daudmannsøyra (*n*=8), which are new in this study, all samples were previously used to generate microsatellite data in a study by Peeters *et al*. (2020).

### DNA extraction, library building, and sequencing

DNA was extracted from ear tissue for the eight samples from Daudmannsøyra using a Qiagen (Hilden, Germany) DNeasy Blood & Tissue extraction kit according to the manufacturer’s instructions except for the addition RNase A (details in SI 1). DNA extraction for all other samples (*n* = 92) is described in Peeters et al. (2020). Genomic library building was performed for all samples based on the method described in (Carøe *et al*. 2018), and 90 of these were then sequenced to a target depth of 2-3x (see Figure S1 and Table S1 for details). These sequencing data were combined with data from deep-sequencing of the remaining ten samples.

### Bioinformatic processing and genotype likelihood calculation

We used Paleomix version 1.2.13.4 (Schubert *et al*. 2014) to map demultiplexed sequence reads to the caribou reference genome assembled from a North American male (Taylor *et al*. 2019). This reference, while more phylogenetically divergent from the Svalbard reindeer, is more contiguous (N_50_ = 11.765 Mbp) than the Mongolian reindeer reference (N_50_ = 0.94 Mbp; (Li *et al*. 2017)) and more suitable for RoH inbreeding-type analyses. Adapters were trimmed with adapterremoval version 2 (Schubert *et al*. 2016) and the BWA aligner program version 0.7.15 was used with the MEM algorithm (Li 2013) without filtering for mapping quality.

To account for the uncertainty in calling genotypes from low-depth sequencing data, we utilised ANGSD v0.93 (Korneliussen *et al*. 2014) to generate genotype likelihood data for each individual, and these, rather than explicitly called genotypes, were used in downstream analyses. Genotype likelihood files were generated in beagle format inferring allele frequencies with fixed major and minor alleles using the command-line arguments *-doGlf 2* (admixture analyses) or *-doGlf3* (inbreeding analyses), *-doMajorMinor 1*, and *-doMaf 1*. Variants were called with a p-value threshold of 1e^-6^ (*-SNP_pval 1e-6*) only at sites for which there was sequence data in at least 50 individuals (*-minInd 50*). Reads with mapping quality <30 and base quality <20, and those with multiple mapping hits, were filtered out using *-minMapQ 30*, *-minQ 20*, and *-uniqueOnly 1*, and low-quality reads were removed with *-remove_bads 1*. Scaffolds mapped to bovine sex chromosomes by Taylor *et al*. (2019) were removed. To reduce any issues related to paralogs or mapping errors we filtered out sites that had average coverage greater than twice or less than ⅓ of the genome-wide average in a sub-sample of 10 individuals with equal (3x) coverage. We used the *-C 50* parameter to adjust map quality for reads with a large number of mismatches to the reference genome, and the extended baq model to adjust quality scores around indels (*-baq 2*).

### Mitochondrial genome analysis

To analyse mtDNA haplotype diversity, we mapped our sequence data to a 16,357 bp reindeer mtDNA reference assembly (Ju *et al*. 2016). Then, we used the GATK 4.1.8.1 HaplotypeCaller (Depristo *et al*. 2011) to identify SNPs and call mtDNA haplotypes. Four samples were excluded from the analysis due to low coverage (<20x). We specified haploid calls (-ploidy 1), only used reads with a minimum mapping quality of 25 (--minimum-mapping-quality 25) and specified a confidence threshold of 30 for variant calling (-stand-call-conf 30). We then converted the haplotype calls of variable sites to FASTA sequences for each individual. To investigate mitochondrial genetic structure and haplotype diversity, we used pegas v1.1 (Paradis 2010) to construct a median-joining haplotype network based on the raw number of nucleotide differences between sequences. We also used pegas to calculate population haplotype richness rarefied to a sample size of 5 using the Hurlbert (1971) rarefaction method.

### Ancestry and admixture analyses

We used the maximum likelihood based clustering analysis software package NGSadmix (Skotte & Albrechtsen 2013) to infer population structure and identify admixture between populations using genotype likelihood data. For admixture analysis, we excluded samples with sequencing depth <0.1x (*n* = 4) and removed two out of three closely related individuals in the reintroduced Daudmannsøyra population identified using NgsRelate (Hanghøj *et al*. 2019), because closely related individuals can bias admixture results (Garcia-Erill & Albrechtsen 2020). We LD pruned the genotype likelihood file based on called genotypes with PLINK v 1.9 (Chang *et al*. 2015) using *--indep-pairwise 50 5 0.3* to specify a window size of 50, step size of 5, and a r^2^ threshold of 0.3. Admixture models were run for the number of genetic clusters (*K*) ranging from 2 – 10, with 10 replicates of each. Only sites with a minimum minor allele frequency greater than 0.02 (using *-minMaf 0.02*) and that had data in at least half (46) the 92 individuals in the analysis (using *–minInd 46*) were included in the analysis. We ran admixture analyses on the full dataset including all populations (467,146 sites), and also a separate analysis on a subset including only the reintroduction source, reintroduced, and Southern Svalbard populations (427,643 sites). For each value of *K*, the replicate with the highest likelihood was selected. We calculated Δ*K* (Evanno *et al*. 2005) using CLUMPAK (Kopelman *et al*. 2015), however uneven sampling of ancestral populations and strong genetic drift can bias rule-based model selection (Garcia-Erill & Albrechtsen 2020). Therefore we considered all *K* models and examined the correlation of residuals using EVALadmix (Garcia-Erill & Albrechtsen 2020) to evaluate and interpret results instead of relying solely on a rule-based model selection procedure.

### Principal component analysis

We conducted Principal component analysis (PCA) using the software package PCAngsd (Meisner & Albrechtsen 2018) to estimate a genetic covariance matrix using individual allele frequencies based on the same genotype likelihood data used in the admixture analysis, then computed eigenvectors and eigenvalues using the *eigen* function in R 3.6 (R Core Team 2019). To visualise the data, we plotted the first four PC axes, and used ggplot2 (Wickham 2016) to calculate 95% CI ellipses of the mean principal component coordinates of each natural population and for Reintroduction 1 and 2. We repeated this analysis using only the individuals from the Adventdalen, Southern Spitsbergen, and reintroduced populations to characterise fine-scale population structure.

### F_ST_ analysis

We quantified population differentiation by estimating pairwise F_ST_ between each population using RealSFS in ANGSD v0.93 based on 2D (pairwise) population site frequency spectra (SFS) including all samples. First, we generated unfolded per-site allele frequencies (SAF) for each population using the *-dosaf 1* argument in ANGSD with the same parameters and site filtering as when generating genotype likelihoods, and specified the reference as the ancestral genome. Then, with the realSFS module in ANGSD, we used the unfolded SAF to generate folded 2D SFS priors for each pair of populations using *-fold 1* since no ancestral states were available to polarise the ancestral/derived alleles. We then input the unfolded SAFs and the folded 2D SFS prior to realSFS to estimate per-site and global F_ST_, specifying the Hudson estimation method which is more suitable for smaller sample sizes (Bhatia *et al*. 2013) using -*whichFST 1*. Finally, we used the realSFS *fst stat* function to calculate the weighted global F_ST_ for each population pair.

### Heterozygosity

We estimated genome-wide heterozygosity for each individual with coverage >2.5x using realSFS in ANGSD v0.93 based on the folded site frequency spectrum of each individual (Korneliussen *et al*. 2014). We used the same site filtering and parameters as for the genotype likelihoods described above, however since coverage can bias heterozygosity estimates in our data (see Fig S11), we downsampled each sample to 2.5x coverage using -DownSample in ANGSD to allow unbiased comparisons between the maximum number of samples. To estimate heterozygosity in ANGSD, we generated a folded SFS from unfolded SAF separately for each individual and divided the number of heterozygous sites by the total number of non-N sites.

### Inbreeding and runs of homozygosity

We used ngsF-HMM (Vieira *et al*. 2016) to identify tracts of individual genomes identical by descent (hereon referred to as RoH), and estimate inbreeding coefficients from genotype likelihoods. This method utilises a hidden Markov model approach to estimate per-site probabilities of being IBD rather than a rule-based method, and can be used with genotyp likelihood data so is more appropriate for use with low-depth WGS data. We excluded scaffolds shorter than 10 Mbp from inbreeding analyses, leaving approximately 56% of the assembled genome (1.235 Gbp) covered by 1,640,852 variable sites on 64 scaffolds after filtering. We also applied stricter filtering than with heterozygosity, restricting this analysis to samples with >2.5x coverage based on reads used by ANGSD on the >10-Mbp scaffolds after all filtering parameters, and then down-sampled each to 2.7x coverage using samtools (Li *et al*. 2009) to allow unbiased RoH comparisons between our samples. We inferred the approximate age of inbreeding (i.e. the number of generations back to the common ancestor that an RoH was inherited from) based on RoH lengths, using the equation G = 100/(2rL) where r is the recombination rate and L is the length of RoH in Mbp (Thompson 2013; Kardos *et al*. 2017), assuming a recombination rate similar to red deer (*Cervus elaphus*) of ∼ 1 cM/Mbp (Johnston *et al*. 2017).

## Acknowledgments

We thank Andrew Foote and Ingerid Hagen for their useful comments on a preliminary draft of this manuscript. We are also grateful to several volunteers for providing tissue samples, including Karoline Bælum at the Farmhamna trapping station. This study was primarily financed by the Research Council of Norway via FRIMEDBIO award 276080 to BBH, FRIMEDBIO award 325589 to MDM, and Arctic Field Grant 295908 to HB. The study also received funding and support from the Research Council of Norway Centres of Excellence SFF-III award 223257, the Svalbard Environmental Protection Fund awards 14/137 and 15/105 to BBH, and the Norwegian Polar Institute for field logistics for some of the data sampling. We acknowledge the following sequencing centres and associates for their invaluable services: Novogene (Cambridge, U.K.), the NTNU Genomics Core Facility (Trondheim, Norway) funded by the NTNU Faculty of Medicine and Health Sciences and the Central Norway Regional Health Authority), the National Sequencing Centre (Oslo, Norway), the Science for Life Laboratory, the Knut and Alice Wallenberg Foundation, the National Genomics Infrastructure funded by the Swedish Research Council, and the Uppsala Multidisciplinary Center for Advanced Computational Science for assistance with massively parallel sequencing and access to the UPPMAX computational infrastructure. Some computational analyses were performed on resources provided by the National Infrastructure for High Performance Computing and Data Storage in Norway (UNINETT Sigma2).

## Author contributions

The study was conceived by BBH, HJ, HB, MDM, and VCB. MDM, BBH, HJ, LEL, and LD provided funding for the study. ÅØP, MLM, BBH and BP collected most samples. HB, VCB, BP, and MDM performed laboratory and bioinformatic analyses. HB wrote the manuscript with contributions from all authors.

## Data availability

The sequence data generated for this study is available on the European Nucleotide Archive under project accession PRJEB57293.

## References

Aanes, R., Sather, B.-E. & Oritsland, N.A. (2000). Fluctuations of an Introduced Population of Svalbard Reindeer : The Effects of Density Dependence and Climatic Variation. Ecography.

Albon, S.D., Irvine, R.J., Halvorsen, O., Langvatn, R., Loe, L.E., Ropstad, E., et al. (2017). Contrasting effects of summer and winter warming on body mass explain population dynamics in a food-limited Arctic herbivore. Glob. Chang. Biol., 23, 1374–1389.

Allendorf, F.W. (1986). Genetic drift and the loss of alleles versus heterozygosity. Zoo Biol., 5, 181–190.

Andersen, A., Simcox, D.J., Thomas, J.A. & Nash, D.R. (2014). Assessing reintroduction schemes by comparing genetic diversity of reintroduced and source populations: A case study of the globally threatened large blue butterfly (*Maculinea arion*). Biol. Conserv., 175, 34–41.

Armbruster, P. & Reed, D.H. (2005). Inbreeding depression in benign and stressful environments. Heredity, 95, 235–242.

Armstrong, D.P. & Seddon, P.J. (2008). Directions in reintroduction biology. Trends Ecol. Evol., 23, 20– 25.

Bertorelle, G., Raffini, F., Bosse, M., Bortoluzzi, C., Iannucci, A., Trucchi, E., et al. (2022). Genetic load: genomic estimates and applications in non-model animals. Nat. Rev. Genet., 0123456789.

Bhatia, G., Patterson, N., Sankararaman, S. & Price, A.L. (2013). Estimating and interpreting FST: The impact of rare variants. Genome Res., 23, 1514–1521.

Biebach, I. & Keller, L.F. (2010). Inbreeding in reintroduced populations: The effects of early reintroduction history and contemporary processes. Conserv. Genet., 11, 527–538.

Biebach, I. & Keller, L.F. (2012). Genetic variation depends more on admixture than number of founders in reintroduced Alpine ibex populations. Biol. Conserv., 147, 197–203.

Bortoluzzi, C., Bosse, M., Derks, M.F.L., Crooijmans, R.P.M.A., Groenen, M.A.M. & Megens, H.J. (2020). The type of bottleneck matters: Insights into the deleterious variation landscape of small managed populations. Evol. Appl., 13, 330–341.

Bozzuto, C., Biebach, I., Muff, S., Ives, A.R. & Keller, L.F. (2019). Inbreeding reduces long-term growth of Alpine ibex populations. Nature Ecology & Evolution, 3, 1359–1364.

Brekke, P., Bennett, P.M., Santure, A.W. & Ewen, J.G. (2011). High genetic diversity in the remnant island population of hihi and the genetic consequences of re-introduction. Mol. Ecol., 20, 29–45.

Brüniche-Olsen, A., Kellner, K.F., Anderson, C.J. & DeWoody, J.A. (2018). Runs of homozygosity have utility in mammalian conservation and evolutionary studies. Conserv. Genet., 19, 1295–1307.

Carlson, S.M., Cunningham, C.J. & Westley, P.A.H. (2014). Evolutionary rescue in a changing world. Trends Ecol. Evol., 29, 521–530.

Carøe, C., Gopalakrishnan, S., Vinner, L., Mak, S.S.T., Sinding, M.H.S., Samaniego, J.A., et al. (2018). Single-tube library preparation for degraded DNA. Methods Ecol. Evol., 9, 410–419.

Cavill, E.L., Gopalakrishnan, S., Puetz, L.C., Ribeiro, Â.M., Mak, S.S.T., da Fonseca, R.R., et al. (2022). Conservation genomics of the endangered Seychelles Magpie-Robin (*Copsychus sechellarum*): a unique insight into the history of a precious endemic bird. Ibis, 164, 396–410.

Chang, C.C., Chow, C.C., Tellier, L.C.A.M., Vattikuti, S., Purcell, S.M. & Lee, J.J. (2015). Second-generation PLINK: Rising to the challenge of larger and richer datasets. Gigascience, 4, 1–16.

Charlesworth, D. & Willis, J.H. (2009). The genetics of inbreeding depression. Nat. Rev. Genet., 10, 783.

Clegg, S.M., Degnan, S.M., Kikkawa, J., Moritz, C., Estoup, A. & Owens, I.P.F. (2002). Genetic consequences of sequential founder events by an island-colonizing bird. Proc. Natl. Acad. Sci. U. S. A., 99, 8127–8132.

Côté, S.D., Dallas, J.F., Marshall, F., Irvine, R.J., Langvatn, R. & Albon, S.D. (2002). Microsatellite DNA evidence for genetic drift and philopatry in Svalbard reindeer. Mol. Ecol., 11, 1923–1930.

Cullingham, C.I. & Moehrenschlager, A. (2013). Temporal Analysis of Genetic Structure to Assess Population Dynamics of Reintroduced Swift Foxes. Conserv. Biol., 27, 1389–1398.

Depristo, M.A., Banks, E., Poplin, R., Garimella, K.V., Maguire, J.R., Hartl, C., et al. (2011). A framework for variation discovery and genotyping using next-generation DNA sequencing data. Nat. Genet., 43, 491–501.

Derocher, A.E., Wiig, Ø. & Bangjord, G. (2000). Predation of Svalbard reindeer by polar bears. Polar Biol., 23, 675–678.

Drauch, A.M. & Rhodes, O.E. (2007). Genetic Evaluation of the Lake Sturgeon Reintroduction Program in the Mississippi and Missouri Rivers. N. Am. J. Fish. Manage., 27, 434–442.

Druet, T. & Gautier, M. (2017). A model-based approach to characterize individual inbreeding at both global and local genomic scales. Mol. Ecol., 26, 5820–5841.

Druet, T., Oleński, K., Flori, L., Bertrand, A.R., Olech, W., Tokarska, M., et al. (2020). Genomic Footprints of Recovery in the European Bison. J. Hered., 194–203.

Duntsch, L., Whibley, A., Brekke, P., Ewen, J.G. & Santure, A.W. (2021). Genomic data of different resolutions reveal consistent inbreeding estimates but contrasting homozygosity landscapes for the threatened Aotearoa New Zealand hihi. Mol. Ecol., 30, 6006–6020.

Evanno, G., Regnaut, S. & Goudet, J. (2005). Detecting the number of clusters of individuals using the software STRUCTURE: a simulation study. Mol. Ecol., 14, 2611–2620.

Fox, C.W. & Reed, D.H. (2011). Inbreeding depression increases with environmental stress: An experimental study and meta-analysis. Evolution, 65, 246–258.

Frankham, R. (2005). Genetics and extinction. Biol. Conserv., 126, 131–140.

Frankham, R. (2010). Challenges and opportunities of genetic approaches to biological conservation. Biol. Conserv., 143, 1919–1927.

Frankham, R., Ballou, J.D., Ralls, K., Eldridge, M., Dubash, M.R., Fenster, C.B., et al. (2017). Genetic Management of Fragmented Animal and Plant Populations. OUP Oxford.

Fuentes-Pardo, A.P. & Ruzzante, D.E. (2017). Whole-genome sequencing approaches for conservation biology: Advantages, limitations and practical recommendations. Mol. Ecol., 26, 5369–5406.

Garcia-Erill, G. & Albrechtsen, A. (2020). Evaluation of model fit of inferred admixture proportions. Mol. Ecol. Resour., 20, 936–949.

Giaimo, S. & Traulsen, A. (2022). Generation time in stage-structured populations under fluctuating environments. Am. Nat., in press.

Gjertz, I. (1995). Reindeer on Prins Karls Forland, Svalbard. Polar Res., 14, 87–88.

Glémin, S. (2003). How are deleterious mutations purged? Drift versus nonrandom mating. Evolution, 57, 2678–2687.

Grossen, C., Biebach, I., Angelone-Alasaad, S., Keller, L.F. & Croll, D. (2018). Population genomics analyses of European ibex species show lower diversity and higher inbreeding in reintroduced populations. Evol. Appl., 11, 123–139.

Grossen, C., Guillaume, F., Keller, L.F. & Croll, D. (2020). Purging of highly deleterious mutations through severe bottlenecks in Alpine ibex. Nat. Commun., 11.

Haanes, H., Røed, K.H., Flagstad & Rosef, O. (2010). Genetic structure in an expanding cervid population after population reduction. Conserv. Genet., 11, 11–20.

Hanghøj, K., Moltke, I., Andersen, P.A., Manica, A. & Korneliussen, T.S. (2019). Fast and accurate relatedness estimation from high-throughput sequencing data in the presence of inbreeding. Gigascience, 8, 1–9.

Hansen, B.B., Aanes, R. & Sæther, B.-E. (2010). Partial seasonal migration in high-arctic Svalbard reindeer (*Rangifer tarandus platyrhynchus*). Can. J. Zool., 88, 1202–1209.

Hansen, B.B., Gamelon, M., Albon, S.D., Lee, A.M., Stien, A., Irvine, R.J., et al. (2019a). More frequent extreme climate events stabilize reindeer population dynamics. Nat. Commun., 10, 1–8.

Hansen, B.B., Pedersen, Å.Ø., Peeters, B., Le Moullec, M., Albon, S.D., Herfindal, I., et al. (2019b). Spatial heterogeneity in climate change effects decouples the long-term dynamics of wild reindeer populations in the high Arctic. Glob. Chang. Biol., 25, 3656–3668.

Hedrick, P.W. & Garcia-Dorado, A. (2016). Understanding Inbreeding Depression, Purging, and Genetic Rescue. Trends Ecol. Evol., 31, 940–952.

Hicks, J.F., Rachlow, J.L., Rhodes, O.E., Williams, C.L. & Waits, L.P. (2007). Reintroduction and Genetic Structure: Rocky Mountain Elk in Yellowstone and the Western States. J. Mammal., 88, 129–138.

Huff, D.D., Miller, L.M. & Vondracek, B. (2010). Patterns of ancestry and genetic diversity in reintroduced populations of the slimy sculpin: Implications for conservation. Conserv. Genet., 11, 2379–2391.

Hundertmark, K.J. & van Daele, L.J. (2010). Founder effect and bottleneck signatures in an introduced, insular population of elk. Conserv. Genet., 11, 139–147.

Hurford, A., Hebblewhite, M. & Lewis, M.A. (2006). A spatially explicit model for an Allee effect: why wolves recolonize so slowly in Greater Yellowstone. Theor. Popul. Biol., 70, 244–254.

Hurlbert, S.H. (1971). The Nonconcept of Species Diversity: A Critique and Alternative Parameters. Ecology, 52, 577–586.

de Jager, D., Glanzmann, B., Möller, M., Hoal, E., van Helden, P., Harper, C., et al. (2021). High diversity, inbreeding and a dynamic Pleistocene demographic history revealed by African buffalo genomes. Sci. Rep., 11, 1–15.

Johansen, B.E., Karlsen, S.R. & Tømmervik, H. (2012). Vegetation mapping of Svalbard utilising Landsat TM/ETM+ data. Polar Rec., 48, 47–63.

Johnston, S.E., Huisman, J., Ellis, P.A. & Pemberton, J.M. (2017). A high-density linkage map reveals sexual dimorphism in recombination landscapes in red deer (*Cervus elaphus*). G3: Genes, Genomes, Genetics, 7, 2859–2870.

Jónsdóttir, I.S. (2005). Terrestrial ecosystems on Svalbard: Heterogeneity, complexity and fragility from an arctic island perspective. Biol. Environ. Proc. R. Ir. Acad., 105B, 155–165.

Ju, Y., Liu, H., Rong, M., Yang, Y., Wei, H., Shao, Y., et al. (2016). Complete mitochondrial genome sequence of Aoluguya reindeer (*Rangifer tarandus*). Mitochondrial DNA. Part A, DNA mapping, sequencing, and analysis, 27, 2261–2262.

Kaeuffer, R., Coltman, D.W., Chapuis, J.L., Pontier, D. & Réale, D. (2007). Unexpected heterozygosity in an island mouflon population founded by a single pair of individuals. Proceedings of the Royal Society B: Biological Sciences, 274, 527–533.

Kardos, M., Armstrong, E.E., Fitzpatrick, S.W., Hauser, S., Hedrick, P.W., Miller, J.M., et al. (2021). The crucial role of genome-wide genetic variation in conservation. Proc. Natl. Acad. Sci. U. S. A., 118.

Kardos, M., Luikart, G. & Allendorf, F.W. (2015). Measuring individual inbreeding in the age of genomics: Marker-based measures are better than pedigrees. Heredity, 115, 63–72.

Kardos, M., Qvarnström, A. & Ellegren, H. (2017). Inferring individual inbreeding and demographic history from segments of identity by descent in Ficedula flycatcher genome sequences. Genetics, 205, 1319–1334.

Kardos, M., Taylor, H.R., Ellegren, H., Luikart, G. & Allendorf, F.W. (2016). Genomics advances the study of inbreeding depression in the wild. Evol. Appl., 9, 1205–1218.

Kekkonen, J., Wikström, M. & Brommer, J.E. (2012). Heterozygosity in an Isolated Population of a Large Mammal Founded by Four Individuals Is Predicted by an Individual-Based Genetic Model. PLoS One, 7, 1–8.

Kohler, J. & Aanes, R. (2004). Effect of winter snow and ground-icing on a Svalbard reindeer population: Results of a simple snowpack model. Arct. Antarct. Alp. Res., 36, 333–341.

Kopelman, N.M., Mayzel, J., Jakobsson, M., Rosenberg, N.A. & Mayrose, I. (2015). Clumpak: A program for identifying clustering modes and packaging population structure inferences across K. Mol. Ecol. Resour., 15, 1179–1191.

Korneliussen, T.S., Albrechtsen, A. & Nielsen, R. (2014). ANGSD: Analysis of Next Generation Sequencing Data. BMC Bioinformatics, 15, 356.

Kutschera, V.E., Poelstra, J.W., Botero-Castro, F., Dussex, N., Gemmell, N.J., Hunt, G.R., et al. (2020). Purifying Selection in Corvids Is Less Efficient on Islands. Mol. Biol. Evol., 37, 469–474.

Kvie, K.S., Heggenes, J., Anderson, D.G., Kholodova, M.V., Sipko, T., Mizin, I., et al. (2016). Colonizing the high arctic: Mitochondrial DNA reveals common origin of Eurasian archipelagic reindeer (*Rangifer tarandus*). PLoS One, 11, 1–15.

Larter, N.C., Sinclair, A.R.E., Ellsworth, T., Nishi, J. & Gates, C.C. (2000). Dynamics of reintroduction in an indigenous large ungulate: the wood bison of northern Canada. Animal Conservation forum, 3, 299–309.

Latch, E.K. & Rhodes, O.E. (2005). The effects of gene flow and population isolation on the genetic structure of reintroduced wild turkey populations: Are genetic signatures of source populations retained? Conserv. Genet., 6, 981–997.

Le Corre, V. & Kremer, A. (1998). Cumulative effects of founding events during colonisation on genetic diversity and differentiation in an island and stepping-stone model. J. Evol. Biol., 11, 495–512.

Le Moullec, M., Pedersen, Å.Ø., Stien, A., Rosvold, J. & Hansen, B.B. (2019). A century of conservation: The ongoing recovery of svalbard reindeer. J. Wildl. Manage., 83, 1676–1686.

Li, H. (2013). Aligning sequence reads, clone sequences and assembly contigs with BWA-MEM. arXiv Preprint, 1–3.

Li, H., Handsaker, B., Wysoker, A., Fennell, T., Ruan, J., Homer, N., et al. (2009). The Sequence Alignment/Map format and SAMtools. Bioinformatics, 25, 2078–2079.

Li, Z., Lin, Z., Ba, H., Chen, L., Yang, Y., Wang, K., et al. (2017). Draft genome of the reindeer *(Rangifer tarandus*). Gigascience, 6, 1–5.

Loe, L.E., Hansen, B.B., Stien, A., Albon, S.D., Bischof, R., Carlsson, A., et al. (2016). Behavioral buffering of extreme weather events in a high-Arctic herbivore. Ecosphere, 7, 1–13.

Loe, L.E., Liston, G.E., Pigeon, G., Barker, K., Horvitz, N., Stien, A., et al. (2021). The neglected season: Warmer autumns counteract harsher winters and promote population growth in Arctic reindeer. Glob. Chang. Biol.

Lønø, O. (1959). Reinen på Svalbard. Norsk Polarinsitutt, Oslo, Norway.

Lynch, M. & Gabriel, W. (1990). Mutation load and the survival of small populations. Evolution, 44, 1725–1737.

MacLeod, I.M., Larkin, D.M., Lewin, H.A., Hayes, B.J. & Goddard, M.E. (2013). Inferring demography from runs of homozygosity in whole-genome sequence, with correction for sequence errors. Mol. Biol. Evol., 30, 2209–2223.

Mathur, S. & DeWoody, J.A. (2021). Genetic load has potential in large populations but is realized in small inbred populations. Evol. Appl., 14, 1540–1557.

Meisner, J. & Albrechtsen, A. (2018). Inferring population structure and admixture proportions in low-depth NGS data. Genetics, 210, 719–731.

Muckenhuber, S., Nilsen, F., Korosov, A. & Sandven, S. (2016). Sea ice cover in Isfjorden and Hornsund, Svalbard (2000-2014) from remote sensing data. Cryosphere, 10, 149–158.

Murphy, S.M., Cox, J.J., Clark, J.D., Augustine, B.C., Hast, J.T., Gibbs, D., et al. (2015). Rapid growth and genetic diversity retention in an isolated reintroduced black bear population in the central appalachians. J. Wildl. Manage., 79, 807–818.

Nei, M., Maruyama, T. & Chakraborty, R. (1975). The Bottleneck Effect and Genetic Variability in Populations. Evolution, 29, 1.

Ochoa, A. & Gibbs, H.L. (2021). Genomic signatures of inbreeding and mutation load in a threatened rattlesnake. Mol. Ecol., 30, 5454–5469.

van Oosterhout, C. (2020). Mutation load is the spectre of species conservation. Nature Ecology and Evolution, 4, 1004–1006.

Paradis, E. (2010). Pegas: An R package for population genetics with an integrated-modular approach. Bioinformatics, 26, 419–420.

Pavlova, O., Gerland, S. & Hopp, H. (2019). Changes in Sea-Ice Extent and Thickness in Kongsfjorden, Svalbard (2003–2016). In: The Ecosystem of Kongsfjorden, Svalbard. Advances in Polar Ecology, vol 2. Springer, Cham, pp. 105–136.

Peeters, B., Le Moullec, M., Raeymaekers, J.A.M., Marquez, J.F., Røed, K.H., Pedersen, Å., et al. (2020). Sea ice loss increases genetic isolation in a high Arctic ungulate metapopulation. Glob. Chang. Biol., 26, 2028–2041.

Peeters, B., Pedersen, Å.Ø., Loe, L.E., Isaksen, K., Veiberg, V., Stien, A., et al. (2019). Spatiotemporal patterns of rain-on-snow and basal ice in high Arctic Svalbard: Detection of a climate-cryosphere regime shift. Environ. Res. Lett., 14.

Pruett, C.L. & Winker, K. (2005). Northwestern song sparrow populations show genetic effects of sequential colonization. Mol. Ecol., 14, 1421–1434.

R Core Team. (2019). R: A Language and Environment for Statistical Computing.

Robinson, J.A., Brown, C., Kim, B.Y., Lohmueller, K.E. & Wayne, R.K. (2018). Purging of Strongly Deleterious Mutations Explains Long-Term Persistence and Absence of Inbreeding Depression in Island Foxes. Curr. Biol., 28, 3487–3494.e4.

Sánchez-Barreiro, F., Gopalakrishnan, S., Ramos-Madrigal, J., Westbury, M.V., de Manuel, M., Margaryan, A., et al. (2021). Historical population declines prompted significant genomic erosion in the northern and southern white rhinoceros (*Ceratotherium simum*). Mol. Ecol., 30, 6355–6369.

Sasmal, I., Jenks, J.A., Waits, L.P., Gonda, M.G., Schroeder, G.M. & Datta, S. (2013). Genetic diversity in a reintroduced swift fox population. Conserv. Genet., 14, 93–102.

Schubert, M., Ermini, L., Sarkissian, C.D., Jónsson, H., Ginolhac, A., Schaefer, R., et al. (2014). Characterization of ancient and modern genomes by SNP detection and phylogenomic and metagenomic analysis using PALEOMIX. Nat. Protoc., 9, 1056–1082.

Schubert, M., Lindgreen, S. & Orlando, L. (2016). AdapterRemoval v2: Rapid adapter trimming, identification, and read merging. BMC Res. Notes, 9, 1–7.

Scott, T.A., Wehtje, W. & Wehtje, M. (2001). The need for strategic planning in passive restoration of wildlife populations. Restor. Ecol., 9, 262–271.

Seddon, P.J., Griffiths, C.J., Soorae, P.S. & Armstrong, D.P. (2014). Reversing defaunation: Restoring species in a changing world. Science, 345, 406–412.

Skotte, L. & Albrechtsen, A. (2013). Estimating Individual Admixture Proportions from. Genetics, 195, 693–702.

Stempniewicz, L., Kulaszewicz, I. & Aars, J. (2021). Yes, they can: polar bears Ursus maritimus successfully hunt Svalbard reindeer Rangifer tarandus platyrhynchus. Polar Biol., 44, 2199–2206.

Stien, A., Loel, L.E., Mysterud, A., Severinsen, T., Kohler, J. & Langvatn, R. (2010). Icing events trigger range displacement in a high-arctic ungulate. Ecology, 91, 915–920.

Stoffel, M.A., Johnston, S.E., Pilkington, J.G. & Pemberton, J.M. (2021). Mutation load decreases with haplotype age in wild Soay sheep. Evolution Letters, 5, 187–195.

Supple, M.A. & Shapiro, B. (2018). Conservation of biodiversity in the genomics era. Genome Biol., 19, 1–12.

Szpiech, Z.A., Xu, J., Pemberton, T.J., Peng, W., Zöllner, S., Rosenberg, N.A., et al. (2013). Long runs of homozygosity are enriched for deleterious variation. Am. J. Hum. Genet., 93, 90–102.

Taylor, R.S., Horn, R.L., Zhang, X., Brian Golding, G., Manseau, M. & Wilson, P.J. (2019). The Caribou (*Rangifer tarandus*) genome. Genes, 10, 4–9.

Taylor, S.S. & Jamieson, I.G. (2008). No evidence for loss of genetic variation following sequential translocations in extant populations of a genetically depauperate species. Mol. Ecol., 17, 545–556.

Thompson, E.A. (2013). Identity by descent: Variation in meiosis, across genomes, and in populations. Genetics, 194, 301–326.

Urbański, J.A. & Litwicka, D. (2022). The decline of Svalbard land-fast sea ice extent as a result of climate change. Oceanologia, 64, 535–545.

Vieira, F.G., Albrechtsen, A. & Nielsen, R. (2016). Estimating IBD tracts from low coverage NGS data. Bioinformatics, 32, 2096–2102.

Vucetich, J.A. & Waite, T.A. (2000). Is one migrant per generation sufficient for the genetic management of fluctuating populations? Anim. Conserv., 3, 261–266.

Wang, J., Hill, W.G., Charlesworth, D. & Charlesworth, B. (1999). Dynamics of inbreeding depression due to deleterious mutations in small populations: Mutation parameters and inbreeding rate. Genet. Res., 74, 165–178.

Weeks, A.R., Sgro, C.M., Young, A.G., Frankham, R., Mitchell, N.J., Miller, K.A., et al. (2011). Assessing the benefits and risks of translocations in changing environments: A genetic perspective. Evol. Appl., 4, 709–725.

Weldenegodguad, M., Pokharel, K., Ming, Y., Honkatukia, M., Peippo, J., Reilas, T., et al. (2020). Genome sequence and comparative analysis of reindeer (*Rangifer tarandus*) in northern Eurasia. Sci. Rep., 10, 1–14.

White, D.J., Watts, C., Allwood, J., Prada, D., Stringer, I., Thornburrow, D., et al. (2017). Population history and genetic bottlenecks in translocated Cook Strait giant weta, Deinacrida rugosa: recommendations for future conservation management. Conserv. Genet., 18, 411–422.

Whitlock, M.C., Ingvarsson, P.K. & Hatfield, T. (2000). Local drift load and the heterosis of interconnected populations. Heredity, 84, 452–457.

Wickham, H. (2016). ggplot2: Elegant Graphics for Data Analysis. Springer-Verlag New York.

Wilder, A.P., Dudchenko, O., Curry, C., Korody, M., Turbek, S.P., Daly, M., et al. (2022). A Chromosome-Length Reference Genome for the Endangered Pacific Pocket Mouse Reveals Recent Inbreeding in a Historically Large Population. Genome Biol. Evol., 14.

Williams, B.W. & Scribner, K.T. (2010). Effects of multiple founder populations on spatial genetic structure of reintroduced American martens. Mol. Ecol., 19, 227–240.

Williams, C.E., Serfass, T.L., Cogan, R. & Rhodes, O.E., Jr. (2002). Microsatellite variation in the reintroduced Pennsylvania elk herd. Mol. Ecol., 11, 1299–1310.

Williams, R.N., Rhodes, O.E. & Serfass, T.L. (2000). Assessment of Genetic Variance Among Source and Reintroduced Fisher Populations. J. Mammal., 81, 895–907.

Willis, K. & Willis, R.E. (2010). How many founders, how large a population? Zoo Biol., 29, 638–646.

Willi, Y., Griffin, P. & Van Buskirk, J. (2013). Drift load in populations of small size and low density. Heredity, 110, 296–302.

Wisely, S.M., Santymire, R.M., Livieri, T.M., Mueting, S.A. & Howard, J. (2008). Genotypic and phenotypic consequences of reintroduction history in the black-footed ferret (*Mustela nigripes*). Conserv. Genet., 9, 389–399.

Wright, D.J., Spurgin, L.G., Collar, N.J., Komdeur, J., Burke, T. & Richardson, D.S. (2014). The impact of translocations on neutral and functional genetic diversity within and among populations of the Seychelles warbler. Mol. Ecol., 23, 2165–2177.

