## Supplementary Information for "Contrasting genomic consequences of anthropogenic reintroduction and natural recolonisation in high-arctic wild reindeer"

[1] Centre for Biodiversity Dynamics, Department of Biology, Norwegian University of Science and Technology (NTNU), Trondheim, Norway; [2] Department of Natural History, NTNU University Museum, Norwegian University of Science and Technology (NTNU), Trondheim, Norway; [3] Department of Terrestrial Biodiversity, Norwegian Institute for Nature Research (NINA), Trondheim, Norway; [4] Norwegian Polar Institute, Research Department, Tromsø, Norway; [5] Centre for Palaeogenetics, Svante Arrhenius väg 20C, SE-106 91 Stockholm, Sweden; [6] Department of Bioinformatics and Genetics, Swedish Museum of Natural History, SE-10405 Stockholm, Sweden; [7] Department of Zoology, Stockholm University, SE-106 91 Stockholm, Sweden; [8] Faculty of Environmental Sciences and Natural Resource Management, Norwegian University of Life Sciences, NO-1432 Aas, Norway; [9] Department of Terrestrial Ecology, Norwegian Institute for Nature Research (NINA), Trondheim, Norway

**SI 1. DNA extraction methods**

A Qiagen (Hilden, Germany) DNeasy Blood & Tissue extraction kit was used to extract DNA from ~20 mg of ear tissue from each of the eight Daudmannsøyra samples, along with an extraction blank. The fur and outer skin layer were removed, and each tissue sample was diced with a scalpel and then placed into a 1.5-ml Eppendorf tube with 180 µL buffer ATL. 20 µL of proteinase K was added to each sample tube and vortexed. Samples were then incubated at 56°C in a Thermoshaker set at 700 RPM for 5.5 hours. 4 µL RNase A (100 mg/ml) was then added to each tube, which were then vortexed and left to incubate at room temp for 2 minutes. Each sample was vortexed after incubation, then 200 µL buffer AL and 200 µL 96% ethanol were added to each sample and immediately vortexed. The mixture was then pipetted into a DNeasy mini spin column placed inside a 2 ml collection tube and centrifuged at 8000 RPM for one minute. Mini spin columns were then placed in a new 2 ml collection tube, 500 µL buffer AW1 added, and centrifuged at 8000 RPM for one minute. Mini spin columns were again placed in a new 2 ml collection tube, 500 µL buffer AW2 added, and centrifuged for three minutes at 14000 RPM. Finally, the mini spin columns were then placed in 1.5 ml centrifuge tubes and 200 µL buffer AE was pipetted onto the DNeasy membrane, and incubated for one minute. The samples were then centrifuged at 8000 RPM for one minute to elute. The DNA concentration was quantified for all using a Thermo Fisher Scientific Qubit 2.0 fluorometer (Indiana, USA).

### SI 2. Library building and sequencing methods

Genomic library building was performed according to the BEST (Blunt End Single Tube) protocol presented in Carøe et al. (2018). Samples were pipetted into a 96-well plate and diluted with EB buffer if necessary, so all wells contain 60 µL volume containing 500 – 5000 ng DNA. A Covaris ME220 focused ultrasonicator was used for DNA fragmentation with a target mean fragment length of 400 bp.

*Blunt end repair.* 8 µL of end-repair master mix (0.4 µL T4 DNA polymerase at 0.03 U/µL concentration, 1 µL T4 polynucleotide kinase at 0.25 U/µL, 0.4 µL dNTPs at 0.25 mM each, 4 µL T4 NDA ligase buffer (NEB) at 1x concentration, and 2.2 µL reaction enhancer) was added to 32 µL of extract from each sample in a 96 well plate and incubated at 20°C for 30 minutes followed by 65°C for 30 minutes and cooled to 4°C.

*Adapter ligation.* 2 µL adapter solution (20 µM) was added and samples were vortexed. 8 µL ligase master mix (1 µL T4 DNA ligase buffer (1x concentration), 6 µL PEG-4000 (6.25%), 1 µL T4 DNA ligase (8 U/µL)). Samples were incubated at 20°C for 30 minutes followed by 65 °C for 10 minutes and cooled to 4 °C.

*Adapter fill-in.* 10 µL of fill-in master mix (2 µL Isothermal amp. Buffer (0.33x concentration), 0.8 µL dNTPs (0.33mM), 1.6 µL Bst 2.0 Warmstart pol. (0.21 U/µL), 5.6 µL $H_2O$) was then added to each sample which were incubated at 65°C for 15 minutes followed by 80°C for 15 minutes and then cooled to 4 °C.

*Library purification.* 100 µL SPRI beads were added to each sample, incubated for five minutes at room temperature, then the plate was placed on a magnetic rack and supernatant was removed once bead pellets had formed. Each sample was washed twice with 200 µL 80% ethanol. The plate was removed from the magnetic rack and 33 µL EBT buffer was used to elute the DNA. The plate was sealed with aluminum foil and incubated at 37 °C for 10 minutes, and then returned to the magnetic rack where the DNA containing supernatant was transferred to a new well plate after bead pellets had formed.

*Indexing PCR and sequencing.* Each sample was indexed with a unique combination of F and R index primers. For each sample, 10 µL of library template and a unique combination of F and R index primers (2 µL of each at 0.2 µM) was added to 96 µL index PCR master mix (0.8 µL dNTPs (0.2 nM), 1 µL AmpliTaq Gold polymerase (0.5 U/µL), 10 µL AmpliTaq Gold buffer (1x conc), 10 μL $MgCl_2$ (2.5 mM), 2 μL BSA (0.4 mg/ml), 62.2 μL $H_2O$), resulting in a total volume of 100μL for each library. All samples were given nine cycles of PCR during indexing. PCR consisted of holding libraries at 95°C for 10 minutes, followed by nine cycles of 95°C for 30 seconds, 60°C for one minute, and 72°C for 45 seconds. After the final cycle, libraries were held at 72°C for five minutes and then cooled to 4°C. 100 μL of SPRI beads (Rohland & Reich, 2012) were mixed with the 100 μL PCR product for each library, which were put onto magnetic racks after incubating at room temperature for five minutes. Libraries were then washed twice with 200 μL 80% ethanol, then removed from the magnetic rack. 33 μL EB buffer was added, libraries were vortexed and incubated at 37°C for 10 minutes and placed back on the magnetic rack. Once beads had reformed, the DNA-containing supernatant was collected and transferred to a new plate. Samples were then run on an Agilent TapeStation 4200 to estimate molarity of samples for pooling. A pooling design was devised based on these molarity values, resulting in samples being combined into eight equimolar pools for sequencing.

### Supplementary tables

**Table S1**. Number of samples from each population sequenced, and number used in population structure analyses (NGSadmix and PCAngsd only), inbreeding analyses ($F_{ROH}$ and runs of homozygosity analysis), heterozygosity estimates, and mitochondrial haplotype analyses.

| **Population** | ***n* Seq** | ***n* Struct** | ***n* Mito** | ***n* Hetero** | ***n* $F_{ROH}$** |
|---|---|---|---|---|---|
| Adventdalen (ADV)[S] | 17 | 17 | 17 | 17 | 17 |
| Brøggerhalvøya (BGR)[R1] | 8 | 8 | 8 | 8 | 7 |
| Sarsøyra (SAR)[R1] | 6 | 6 | 6 | 6 | 5 |
| Kaffiøyra (KAF)[R1] | 9 | 9 | 9 | 9 | 9 |
| Prins Karls Forland (KAF)[R1] | 6 | 5 | 5 | 5 | 4 |
| Daudmannsøyra (DAU)[R2] | 8 | 6 | 8 | 8 | 7 |
| North Isfjorden (NIF)[R2] | 9 | 8 | 8 | 8 | 6 |
| Wijdefjorden (WDF) | 8 | 8 | 8 | 8 | 8 |
| Mitrahalvøya (MTR) | 8 | 7 | 7 | 6 | 5 |
| Southern Spitsbergen (STH) | 8 | 7 | 7 | 6 | 4 |
| Eastern Svalbard (EST) | 10 | 10 | 10 | 9 | 9 |
| North East Land (NE) | 3 | 3 | 3 | 3 | 2 |
| Total | 100 | 94 | 96 | 93 | 83 |

[S] Reintroduction source population; [R1] Reintroduction 1; [R1] Reintroduction 2

**Table S2.** Mitochondrial DNA haplotype and haplogroup counts and rarefied haplotype richness (to a sample size of 5) for all populations and reintroduced populations. Haplogroups are defined as in Fig 4. Haplotype richness was not calculated for the population NE due to its low sample size. ADV: Adventdalen; BGR: Brøggerhalvøya; SAR: Sarsøyra; KAF: Kaffiøyra; PKF: Prins Karls Forland; DAU: Daudmannsøyra; NIF: North Isfjorden; WDF: Wijdefjorden; MTR: Mitrahalvøya; STH: Southern Spitsbergen; EST: Eastern Svalbard; NE: North East Land.

| Population | Sample size | N haplotypes | Rarefied haplotype richness | N haplogroups |
|---|---|---|---|---|
| ADV[S] | 17 | 5 | 2.95 | 3 |
| BGR[R1] | 8 | 3 | 2.25 | 2 |
| PKF[R1] | 5 | 2 | 2.00 | 2 |
| SAR[R1] | 6 | 2 | 2.00 | 2 |
| KAF[R1] | 9 | 4 | 3.29 | 4 |
| Reintro 1 combined | 28 | 5 | 2.79 | 4 |
| DAU[R2] | 8 | 3 | 2.61 | 3 |
| NIF[R2] | 8 | 3 | 2.50 | 2 |
| Reintro 2 combined | 16 | 5 | 2.87 | 4 |
| STH | 7 | 2 | 1.95 | 1 |
| EST | 10 | 2 | 1.50 | 1 |
| NE | 3 | 1 | - | 1 |
| WDF | 8 | 4 | 3.14 | 4 |
| MTR | 7 | 2 | 1.71 | 1 |
| Total | 96 | 16 | | 7 |

[S] Reintroduction source population; [R1] Reintroduction 1; [R1] Reintroduction 2

**Table S3**. Provenance of samples used in this study. See attached .xlsx file

### Supplementary figures

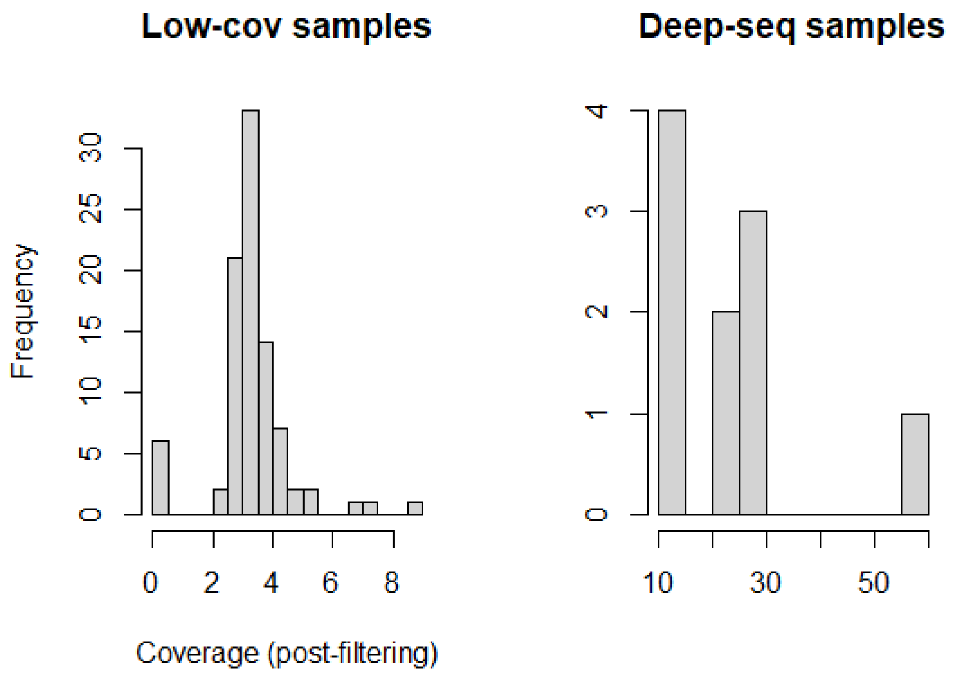


**Figure S1.** Sequencing coverage distribution of sequenced reindeer samples mapped to reindeer and caribou genomes from high coverage and low coverage samples. Coverage calculated after filtering for mapping and base quality.

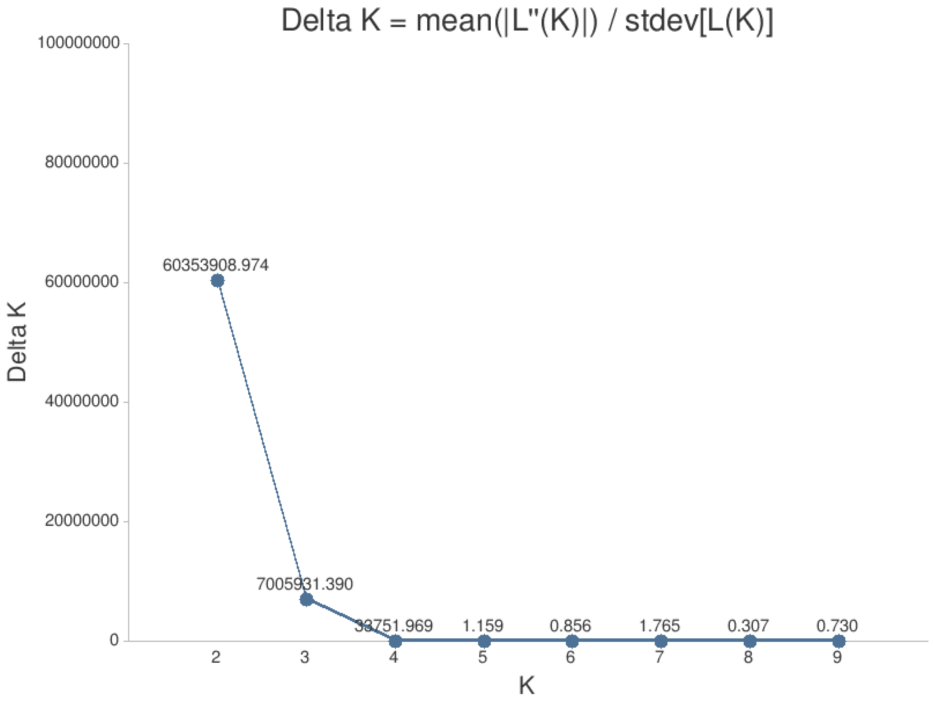


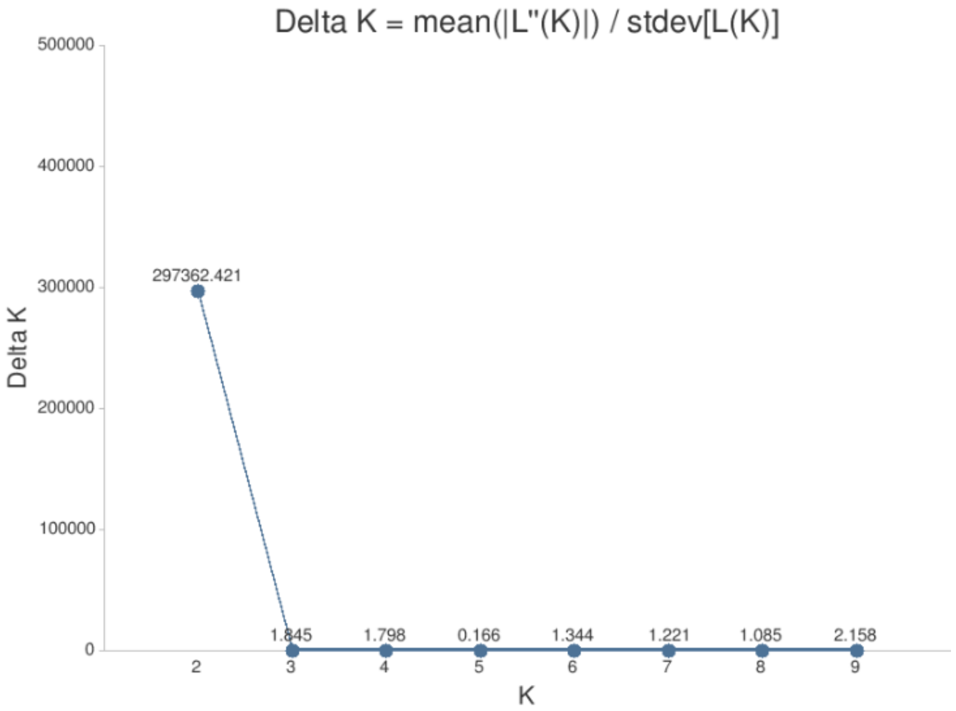


**Figure S2.** Top: Delta *K* values calculated using CLUMPAK for admixture models including the Svalbard-wide dataset (top) and Central Svalbard populations only (bottom).

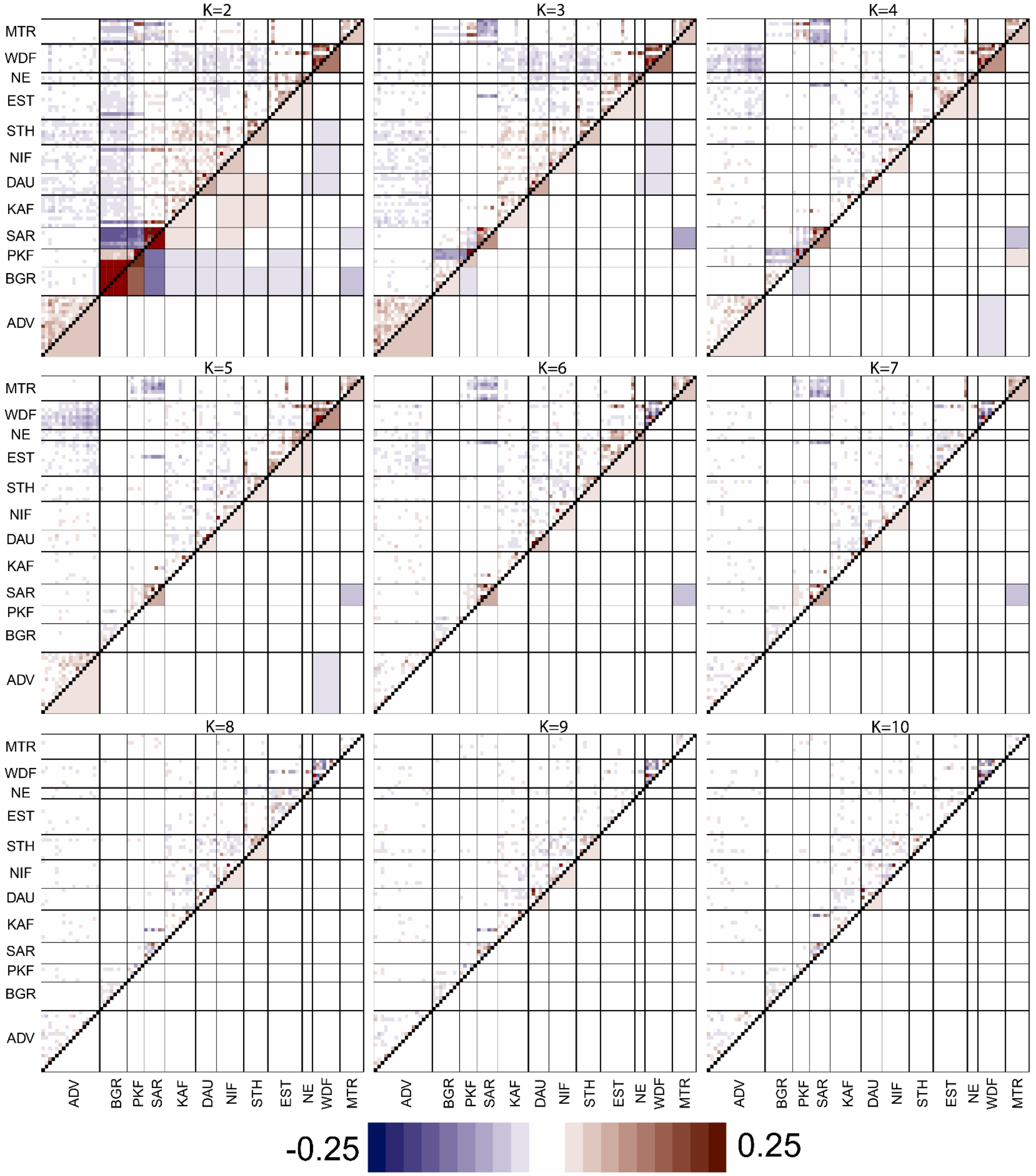


**Figure S3a.** Evaluation of admixture models for the Svalbard-wide dataset, *K*=2 to *K*=10. Colour indicates residual correlation value. Above diagonal shows correlation of residuals between individuals and below diagonal show mean residual correlation within and between populations. ADV: Adventdalen; BGR: Brøggerhalvøya; SAR: Sarsøyra; KAF: Kaffiøyra; PKF: Prins Karls Forland; DAU: Daudmannsøyra; NIF: North Isfjorden; WDF: Wijdefjorden; MTR: Mitrahalvøya; STH: Southern Spitsbergen; EST: Eastern Svalbard; NE: North East Land.

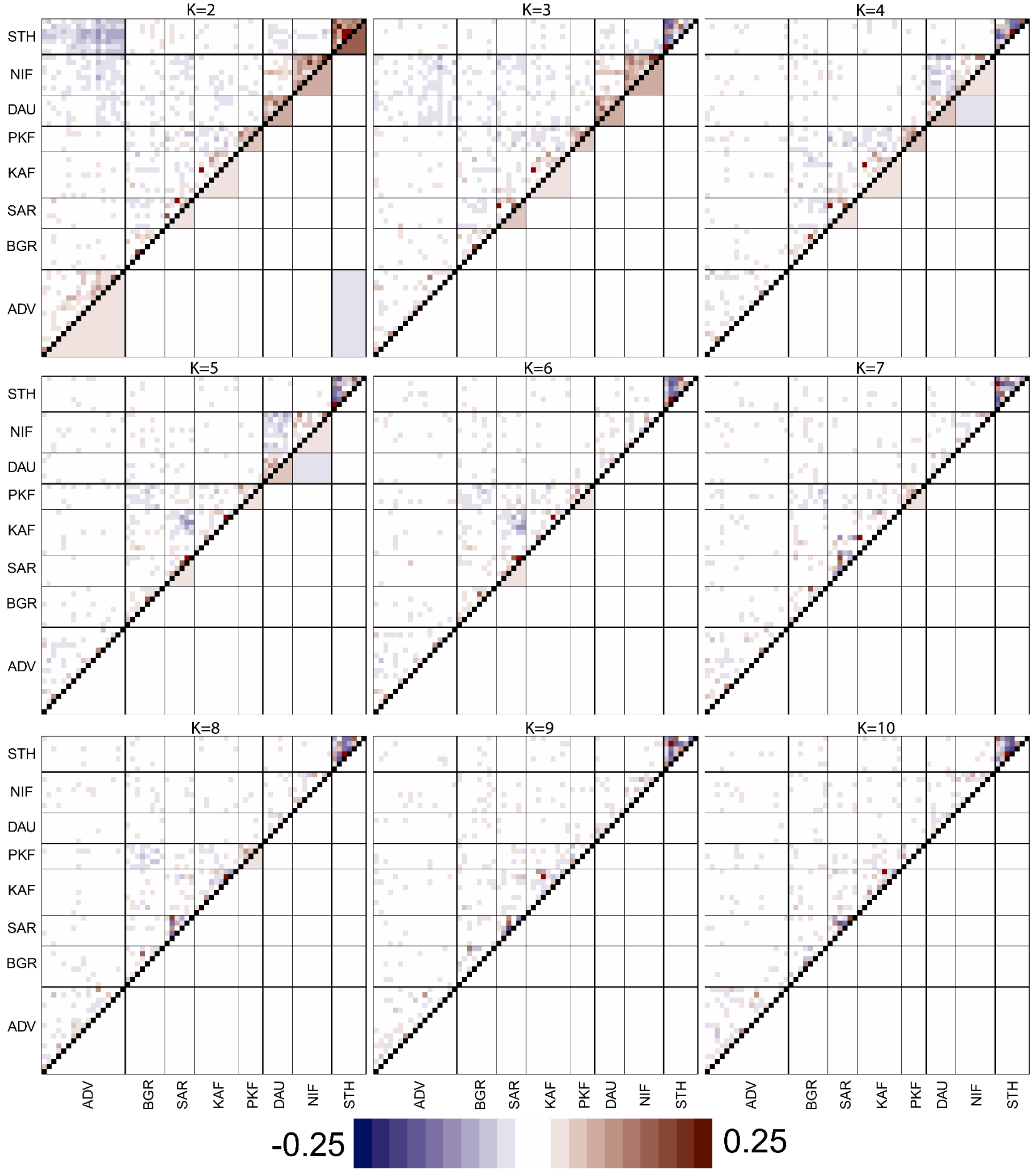


**Figure S3b.** Evaluation of admixture models for the Central Svalbard cluster, *K*=2 to *K*=10. Colour indicates residual correlation value. Above diagonal shows correlation of residuals between individuals and below diagonal show mean residual correlation within and between populations. ADV: Adventdalen; BGR: Brøggerhalvøya; SAR: Sarsøyra; KAF: Kaffiøyra; PKF: Prins Karls Forland; DAU: Daudmannsøyra; NIF: North Isfjorden; STH: Southern Spitsbergen.

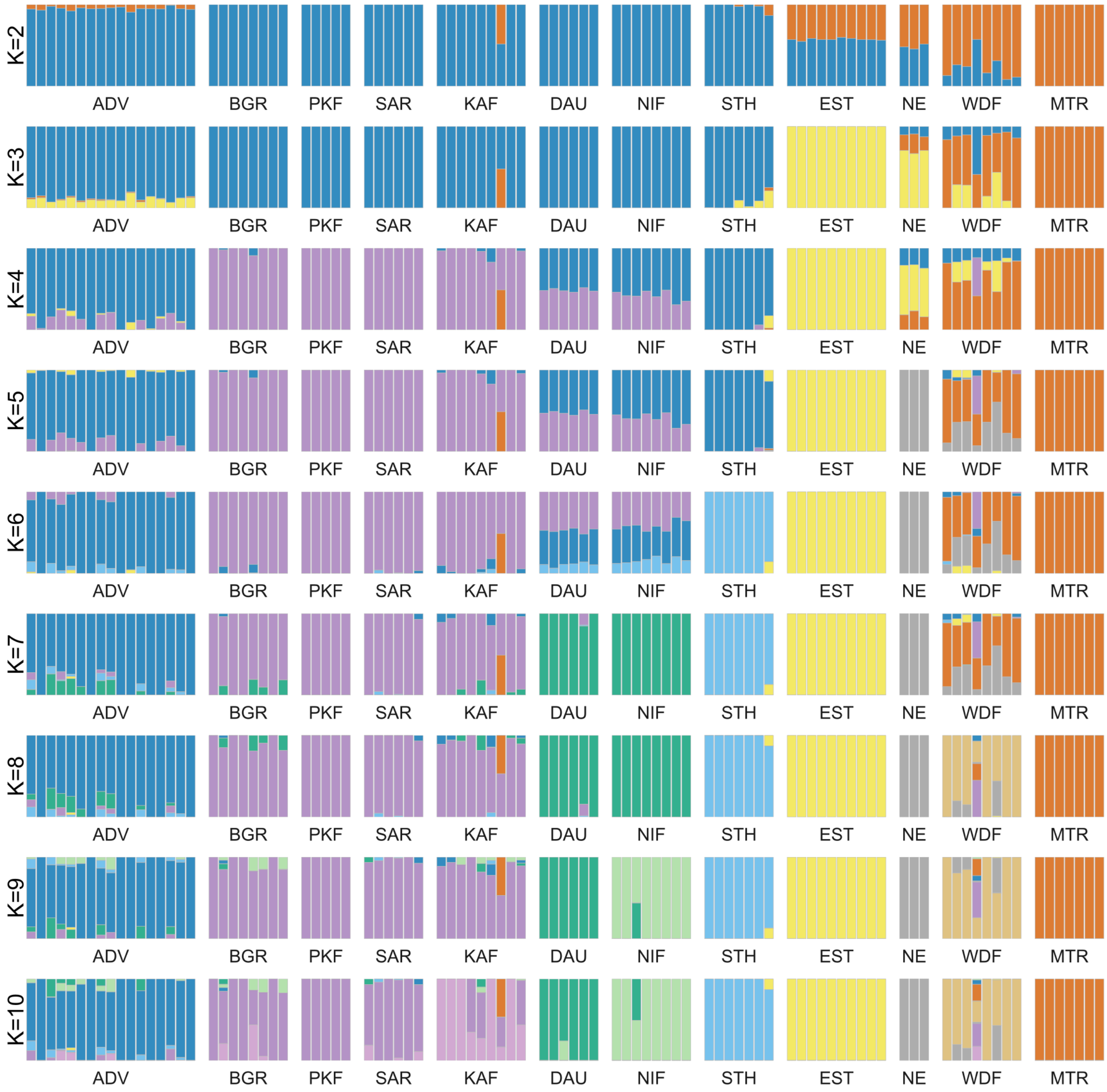


**Figure S4.** Admixture analysis results from NGSadmix models *K*=2 - *K*=10 for the whole Svalbard metapopulation. Vertical bars represent individual reindeer and colours correspond to genetic cluster assignment. ADV: Adventdalen; BGR: Brøggerhalvøya; SAR: Sarsøyra; KAF: Kaffiøyra; PKF: Prins Karls Forland; DAU: Daudmannsøyra; NIF: North Isfjorden; WDF: Wijdefjorden; MTR: Mitrahalvøya; STH: Southern Spitsbergen; EST: Eastern Svalbard; NE: North East Land.

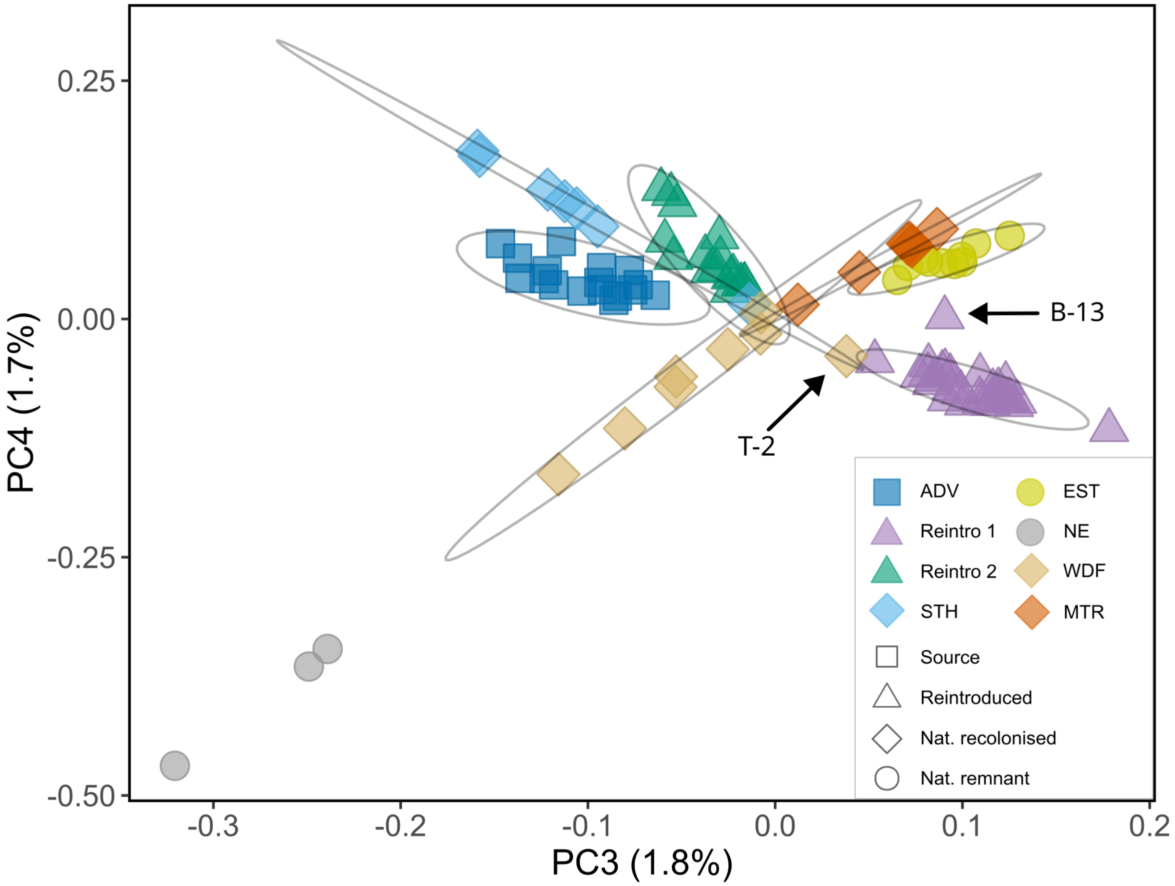


**Figure S5**. Svalbard reindeer genetic PCA PC axes 3 and 4 for the Svalbard-wide metapopulation. 95% CI ellipses for each natural population and each reintroduction group consisting of reintroduced populations grouped according to their expected reintroduction of origin using ggplot2. Two individuals (T-2 and B-13) that represented admixture between strongly differentiated populations were not included in population ellipse calculations. ADV: Adventdalen; BGR: Brøggerhalvøya; SAR: Sarsøyra; KAF: Kaffiøyra; PKF: Prins Karls Forland; DAU: Daudmannsøyra; NIF: North Isfjorden; WDF: Wijdefjorden; MTR: Mitrahalvøya; STH: Southern Spitsbergen; EST: Eastern Svalbard; NE: North East Land.

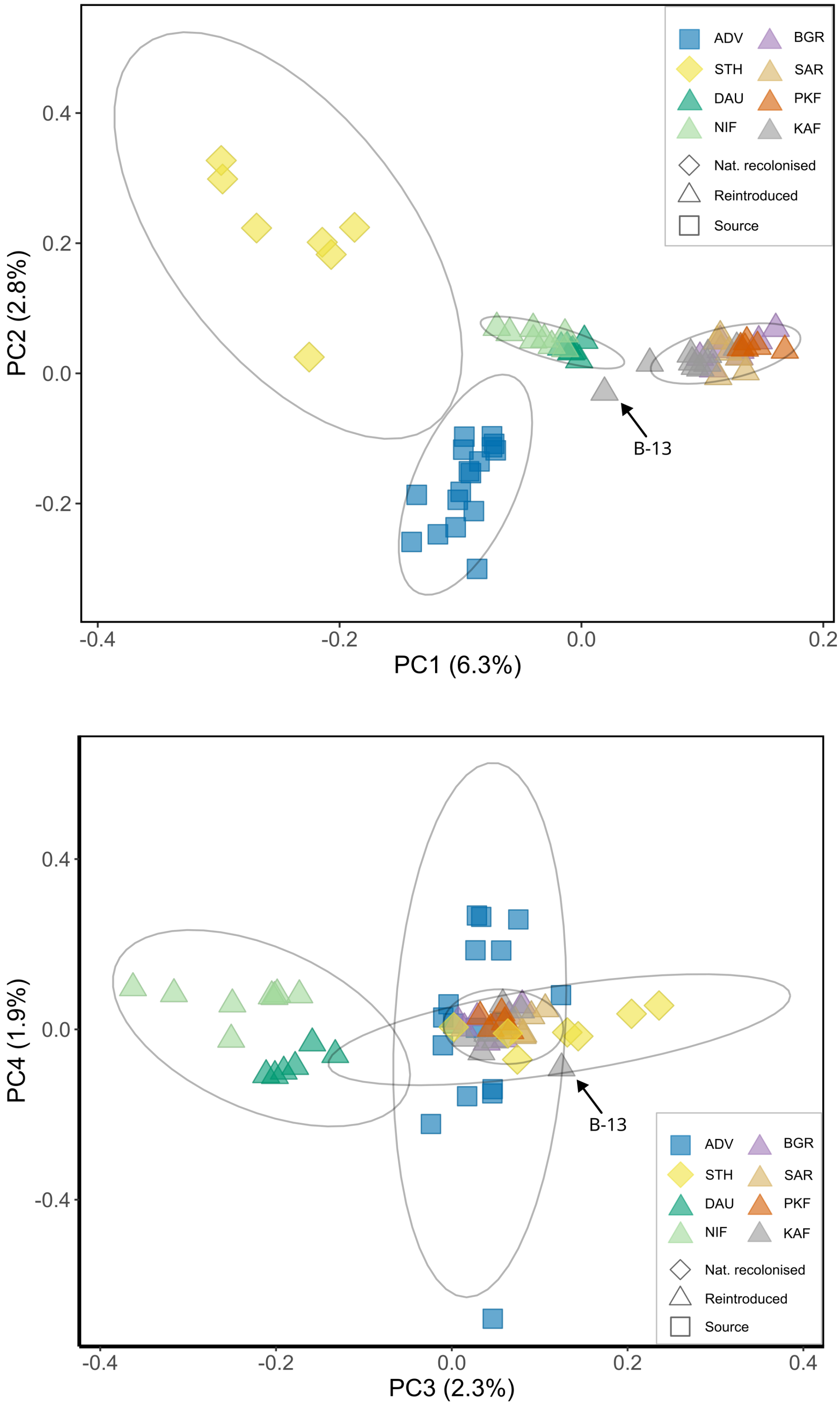


**Figure S6.** Reindeer genetic PCA Principal components 1 and 2 (top) and principal components 3 and 4 (bottom) including only populations from the Central Svalbard group. Ellipses represent the 95% CI of the mean PC coordinates for each natural population and each reintroduction group. One Admixed individual (B-13) in KAF was excluded from ellipse calculations. ADV: Adventdalen; BGR: Brøggerhalvøya; SAR: Sarsøyra; KAF: Kaffiøyra; PKF: Prins Karls Forland; DAU: Daudmannsøyra; NIF: North Isfjorden; STH: Southern Spitsbergen.

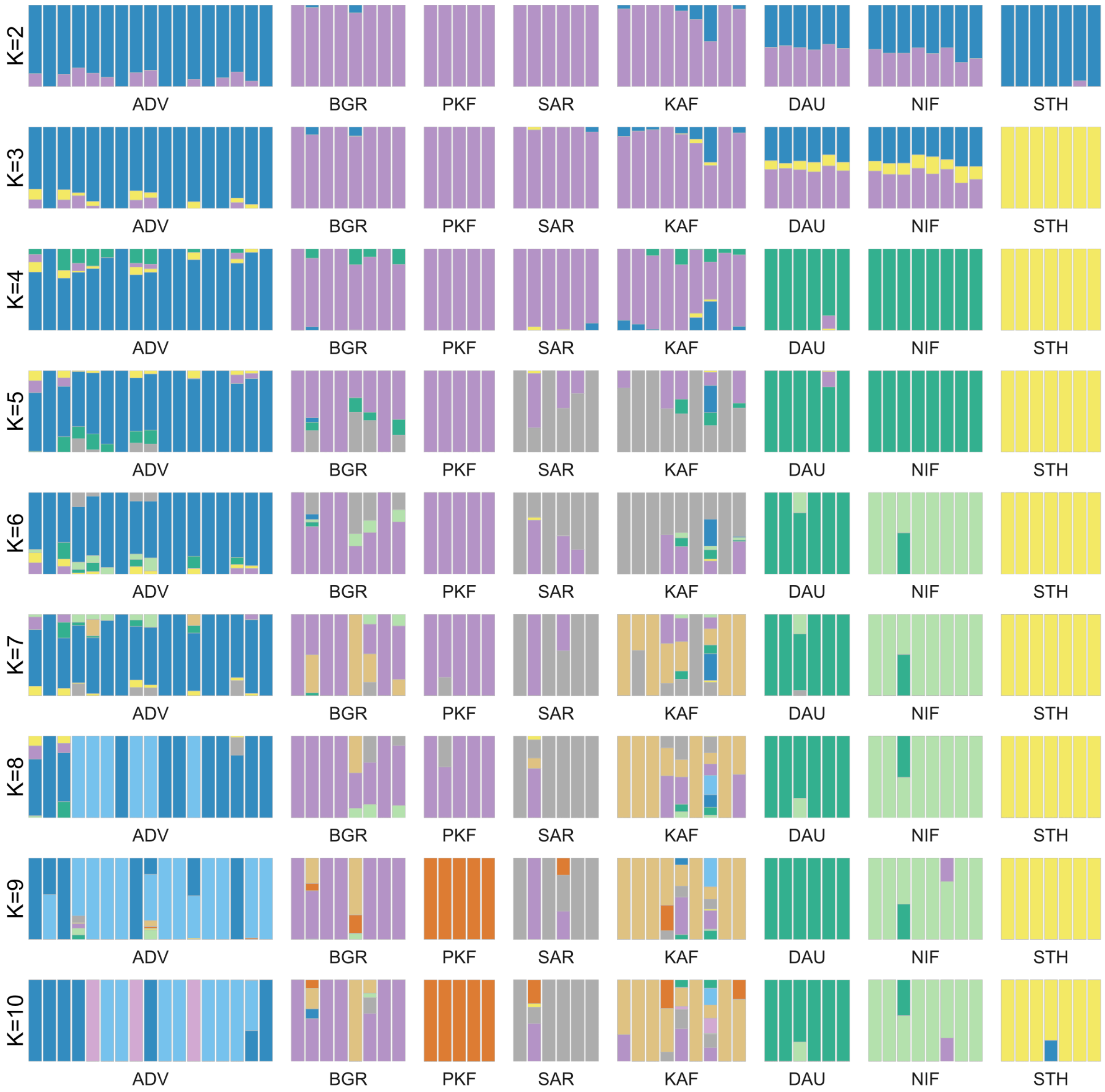


**Figure S7.** Admixture analyses of Svalbard reindeer including only populations from the Central Svalbard group. Vertical bars represent individual reindeer and colours correspond to genetic cluster assignment. ADV: Adventdalen; BGR: Brøggerhalvøya; SAR: Sarsøyra; KAF: Kaffiøyra; PKF: Prins Karls Forland; DAU: Daudmannsøyra; NIF: North Isfjorden; STH: Southern Spitsbergen.

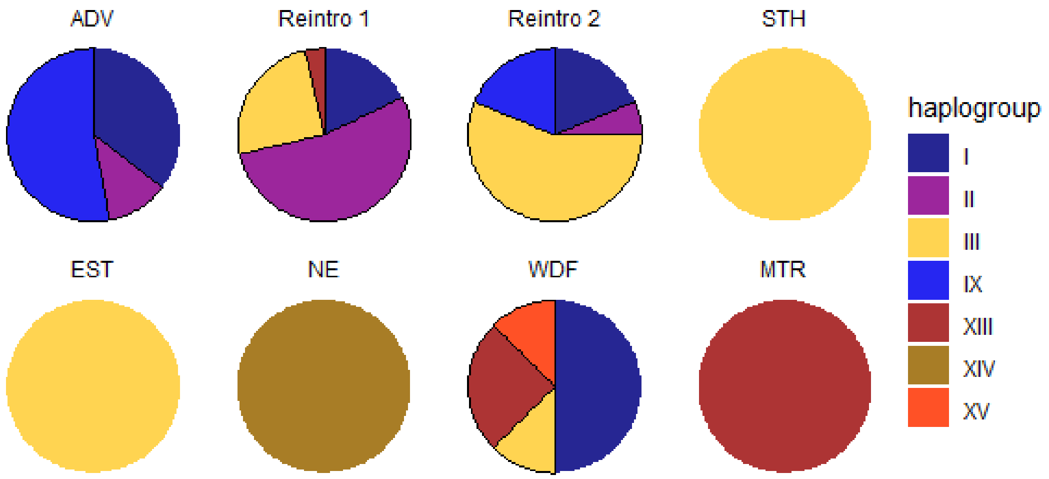


**Figure S8** Mitochondrial haplogroup diversity of each population based on haplotype groupings in Fig. 6. (Reintro 1 (BGR, SAR, KAF, PKF) and Reintro 2 (DAU, NIF) combined, and ADV and STH individually). ADV: Adventdalen; BGR: Brøggerhalvøya; SAR: Sarsøyra; KAF: Kaffiøyra; PKF: Prins Karls Forland; DAU: Daudmannsøyra; NIF: North Isfjorden; WDF: Wijdefjorden; MTR: Mitrahalvøya; STH: Southern Spitsbergen; EST: Eastern Svalbard; NE: North East Land.

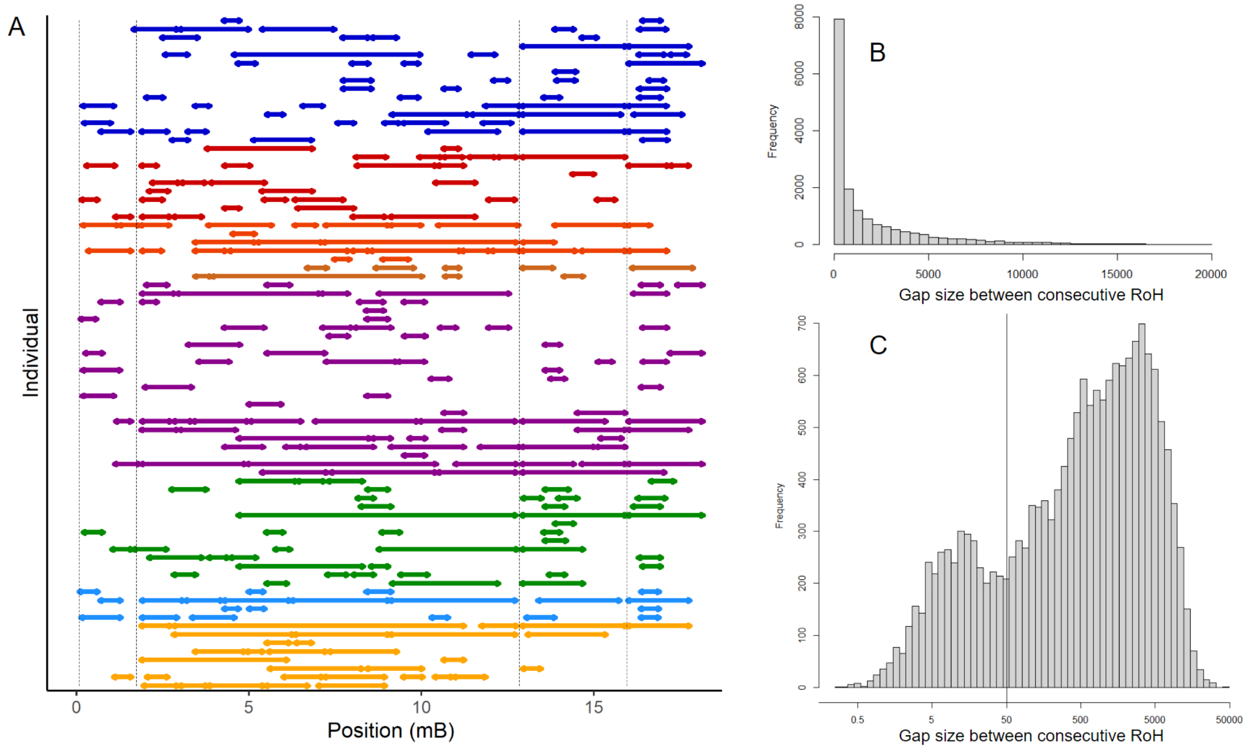


**Figure S9.** Distribution of gap sizes between consecutive RoH. A) RoH plotted for each individual on scaffold Sc247Nf_1653. Coloured segments indicate regions within RoH and arrow endings indicate RoH segment ends. Colours represent separate populations and dashed vertical lines indicate RoH end points shared by all individuals. B) Frequency distribution of gap sizes between consecutive RoH across all individuals (normal scale). C) Frequency distribution of RoH gap sizes on log-scale.

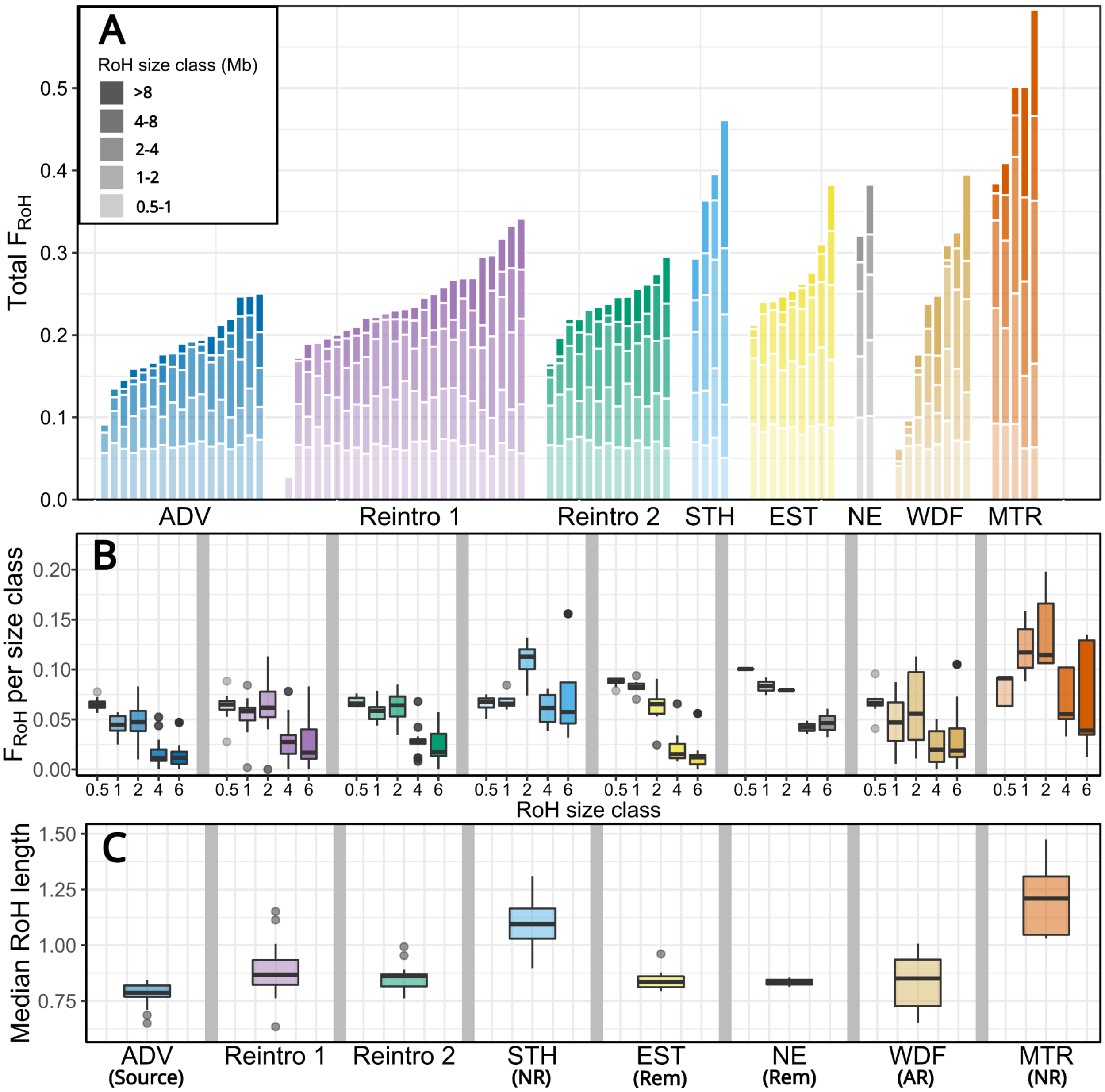


**Figure S10.** RoH results after allowing one 50-Kbp gap within each 1-Mbp RoH. a) Cumulative total $F_{ROH}$ from the five runs of RoH size classes (0.5-1.0 Mbp, 1-2 Mbp, 2-4 Mbp, 4-8 Mbp, and > 8 Mbp) with each bar representing an individual; b) Proportion of individual genomes within RoH of each size classes; c) Median RoH lengths of individuals in each population. NR: Non-admixed naturally recolonised population; AR: Admixed naturally recolonised population; Rem: Natural remnant population. ADV: Adventdalen; BGR: Brøggerhalvøya; SAR: Sarsøyra; KAF: Kaffiøyra; PKF: Prins Karls Forland; DAU: Daudmannsøyra; NIF: North Isfjorden; WDF: Wijdefjorden; MTR: Mitrahalvøya; STH: Southern Spitsbergen; EST: Eastern Svalbard; NE: North East Land.

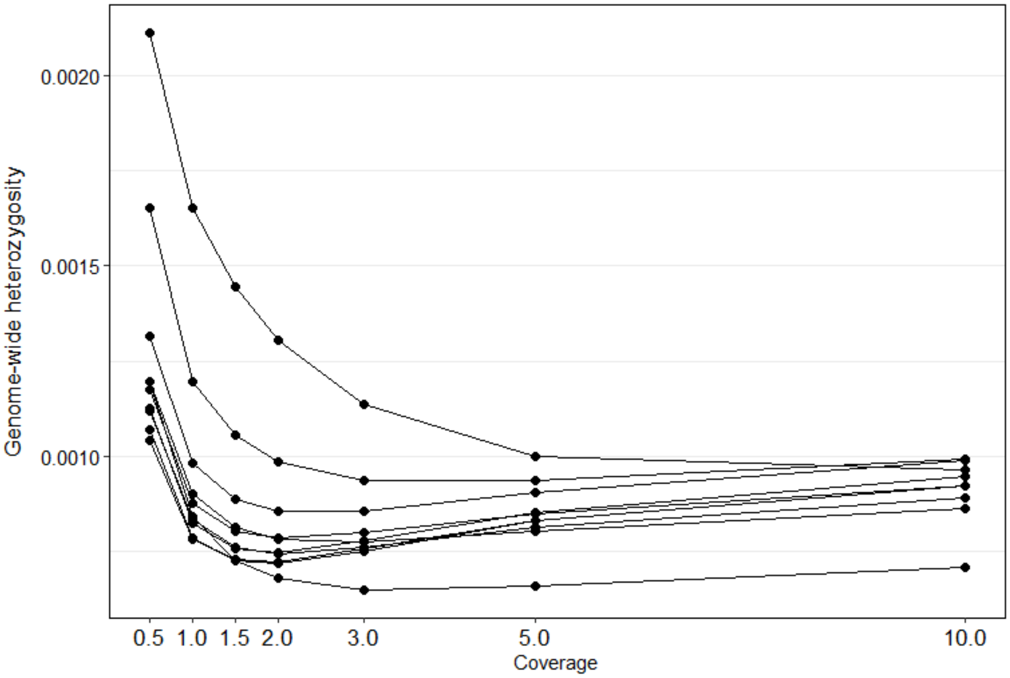


**Figure S11.** Effect of sequencing coverage on heterozygosity based on downsampling 10 individual samples (each line) to 10x, 5x, 3x, 2x, 1.5x, 1x, and 0.5x. Heterozygosity estimates were based on the GL1 genotype likelihood model in ANGSD without the -C50 or -baq 2 parameters and without filtering out sites with high or low average coverage.
